# Mobilizable adipose stromal cells fuel regenerative adipogenesis in injured muscle

**DOI:** 10.1101/2025.09.23.678035

**Authors:** Margaux Labrosse, Maxime Mathieu, Laura Le Pelletier, Virginie Bourlier, Corinne Barreau, Mireille André, Emmanuelle Arnaud, Cédric Moro, Coralie Sengenès, Amandine Girousse, Xavier Contreras

## Abstract

Skeletal muscle regeneration is a highly orchestrated process involving the dynamic interplay of multiple cell types. Among these, fibro-adipogenic progenitors (FAPs), a population of resident mesenchymal stromal cells (MSCs), are essential for creating a supportive microenvironment that promotes satellite cell differentiation and modulates immune responses. Our recent work revealed that adipose stromal cells (ASCs) from subcutaneous adipose tissue (ScAT) infiltrate the injured muscle within the first 24h post-injury, contributing significantly to the regenerative process. Consequently, the FAP population in the regenerating muscle comprises both resident FAPs and infiltrated ASCs.

In the present study, using single cell RNA-seq in a mouse model with trackable KikGR^+^ ASCs and bioinformatics analyses, we identify Limch1^+^/Prg4^+^ ASCs as the primary Mobilizable ASCs (Mob-ASCs) that migrate to and infiltrate the injury site. Notably, this migration is detectable as early as 14 hours post-injury. We demonstrate that these cells are pre-activated within the ScAT, primed to initiate both migratory and regenerative programs. Intriguingly, bioinformatic inference of key activated transcription factors suggested that adipogenesis is also activated in these cells. Leveraging supervised machine learning, we tracked the fate of Mob-ASCs within the regenerating muscle post-injury, where they continue to execute these programs. Importantly, these cells lineage is cued towards a fate of adipogenesis. *In vivo*, we observed transient generation of adipocytes with a peak at 7-9 days post-injury to which infiltrated ASCs contributed. *In vitro*, conditioned media assays further revealed that adipocytes derived from ASCs—but not those from FAPs—enhance myoblasts fusion.

Collectively, our findings establish Limch1^+^/Prg4^+^ ASCs as the Mobilizable ASC population and suggest that their transient adipogenic differentiation is beneficial for muscle regeneration.

## Introduction

Skeletal muscle regeneration is orchestrated by the interplay of various cell types that coordinate the repair and remodeling of damaged tissue^1,2^. While muscle stem cells (MuSCs) are the primary effectors of myofiber regeneration, they rely on the surrounding niche for guidance, support, and modulation of their function. Among the niche components, fibro/adipogenic progenitors (FAPs) have emerged as central players in the muscle repair through their secretion of extracellular matrix (ECM) components, immunoregulatory factors, and trophic signals that modulate MuSCs activity and local inflammation^3–9^. Indeed, inhibition of FAP expansion significantly alters the dynamics of MuSCs, leading to premature MuSCs differentiation, depletion of the early MuSCs pool, and smaller regenerated myofibers, ultimately impairing muscle repair^3,10–15^.

Following acute injury, FAPs are activated and undergo massive expansion around 3 days post-injury through FAP proliferation^3,16–19^. Interestingly, studies have reported a sharp increase in FAP number as early as 24 hours after damage, before proliferation occurs^20^. The origin of this rapid cellular influx has remained elusive, suggesting the possible recruitment of extra muscular mesenchymal stromal cells (MSCs) in the light of our concomitant work demonstrating the possibility of ASCs to egress ScAT in various contexts of tissue stress^21^. Recently, we further demonstrated that adipose stromal cells (ASCs) from subcutaneous adipose tissue (scAT), migrate into the injured muscle and contribute to the FAP compartment playing a supportive role in muscle regeneration^20^. This result reveals an unexpected yet functionally essential dialog between the scAT and skeletal muscle, consistent with our previous observations that scAT releases ASCs upon exposure to inflammatory stimuli^21,22^ or metabolic stress such as high-fat diet^23^, suggesting a broader role for ASCs in responding to diverse environmental cues.

Single-cell analyses in various adipose depots have revealed a functional heterogeneity within ASC populations, uncovering subpopulations with distinct roles^24–26^. Despite the recent discovery that ASCs contribute to muscle regeneration, it remains entirely unknown whether only a specific ASC subpopulation is recruited and functionally engaged during muscle repair. Even more elusive is what these cells actually do once they infiltrate the muscle.

Our study reveals that Progenitor ASCs within the scAT are activated after injury to give rise to Limch1+Prg4+ ASCs that we identified as the subset migrating into the muscle, where it contributes to injury-induced FAP subpopulations. Using integrated bioinformatics, we traced their early activation and defined a unique molecular signature. These ASCs-derived FAPs are less fate-restricted than resident FAPs, retaining greater plasticity and favoring adipogenic differentiation. Strikingly, part of them are capable of transient adipogenesis forming adipocytes whose secretions are notably more pro-myogenic than those from FAPs, supporting myogenic maturation and pointing to a key role for this transient population in muscle regeneration.

## Results

### Specific ASC subpopulations are mobilized upon muscle injury

To identify which ASC subpopulations migrate from scAT to the muscle following injury and to trace their fate, we performed single-cell RNAseq (scRNA-seq). However, ASCs and local muscle-resident FAPs share overlapping markers, so we cannot track ASCs by scRNA-seq into the muscle without an external marker. To overcome this, we used the previously validated KikGR mouse model^20^, in which pieces of KikGR^+^ scAT from a donor was grafted into a wild-type recipient scAT (Fig. 1A). At 7 days post-graft, we induced quadriceps injury using glycerol, or left muscles uninjured. At 1 day post-injury (dpi), we collected both muscle and scAT and isolated ASCs (Sca-1^+^CD31^-^/CD45^-^/Ter119^-^). We then performed scRNA-seq to investigate their cell composition. After quality filtering and doublet removal, we retained 12259 cells (3076 ASCs and 2032 FAPs in the non-injured condition, 3261 ASCsand 3890 FAPs in the injured condition. ASCs and FAPs were first analysed separately to define their subpopulation structure (Supplementary Fig. 1B-C). In ASCs, we identified a total of six subpopulations. Three subpopulations, each segregated by the markers Limch1, Vit+ and Fgf10 correspond respectively to subsets described repeatedly in the literature as Progenitors ASCs (Dpp4^+^/Cd26^+^), Aregs (F3^+^/Cd142^+^) and Preadipocytes (Icam1^+^/Cd54^+^). ^24^. The three other subsets correspond to less characterized Cthrc1^+^, Lgr5^+^ and an injury-induced Mki67^+^ proliferative clusters. FAPs clustered into eight subpopulations enriched in ECM remodeling-related programs, including Hmcn1^+^ (tissue morphogenesis), Hsd11b1^+^(motility/angiogenesis), Pi16^+^ (PI3K/AKT-mediated cell growth signaling), and C7^+^ (fibrogenesis/immunomodulation) as well as injury-specific Fosl1^+^ (chemotaxis/transcriptional activation), Sema3d^+^ and Smim41^+^, and Mki67^+^ (proliferative) populations. To explore potential ASC-to-FAP relationships, we integrated datasets and visualized subpopulation proximity in UMAP space. The resulting UMAP revealed close proximity between three ASC-FAP pairs: ASCs Vit^+^ and FAPs C7^+^, ASCs Limch1^+^ and FAPs Pi16^+^ or Mki67^+^ respectively (Fig. 1B) supporting our hypothesis. To directly identify migrating ASCs within the muscle, we tracked KikGR^+^ cells in FAPs at 1 dpi. We found 577 KikGR^+^ FAPs, corresponding to cells derived from grafted scAT and infiltrated into the injured muscle, predominantly in the injury-specific Fosl1^+^ and Mki67^+^ FAP clusters (Fig. 1C). While these cells represent only graft-derived ASCs, it is likely that endogenous non-fluorescent ASCs also contribute, although they remain unlabelled (Fig. 1C). **Thus, these findings suggest that ASCs contribute to the emergence of FAP subpopulations observed in response to muscle injury but are not sufficient do determine which ASCs are involved.** To identify candidate migrating ASCs, we conducted bioinformatics analyses. We first performed joint clustering of ASCs and FAPs to identify candidate sub-populations of ASCs migrating to muscle upon injury. This revealed three ASC-FAP hybrid clusters: ASC-FAP1 (Vit+ ASCs and C7+/Sema3d+ FAPs), ASC-FAP2 (Limch1+ ASCs and Pi16+ FAPs) and ASC-FAP3 (Mki67+ ASCs and Mki67+ FAPs) matching UMAP proximity results. Indeed, these were the subpopulations already identified as transcriptionally close on the UMAP. Pair wise correlation analysis confirmed transcriptional similarity within these clusters (Fig 1E). ASC-FAP3 is a strong candidate as we found KikGR+ cells in the muscle part of the cluster. However, the 2 other candidates do not have anything in common with KikGR+ cluster. These results are a snapshot at one day post-injury. It is possible that some of these candidates had time to undergo differentiation. Thus, to confirm these candidates, trajectory inference using PAGA Tree was performed on injured samples. This revealed two main paths toward the Fosl1^+^ FAP state (Fig. 1F, Supplementary Fig. 1G). One originated from Limch1^+^ ASCs and bifurcates into two paths: one leading toward other ASC subpopulations, and the other progressing via Pi16^+^ FAPs, marked by the presence of KikGR^+^ cells along the path, and terminating in the Fosl1+ FAP subpopulation, which is also enriched in KikGR^+^ cells. A second trajectory arises from the Mki67^+^ ASCs cluster and also converges on the Fosl1^+^ FAP subpopulation but following a distinct path via the Mki67^+^ FAP subset, which is similarly enriched in KikGR^+^ cells. These results suggest that two ASC subsets may contribute to the same FAP terminal state through distinct differentiation trajectories. Consistently, RNA velocity analysis corroborated the inferred trajectories, confirming a directional flow from Limch1^+^ ASCs and Mki67^+^ ASCs toward terminal FAP states enriched in KikGR^+^ cells (Supplementary Fig. 1H). **These findings establish both a transcriptional and dynamic link between specific ASC subpopulations, Limch1+ and Mki67+ ASCs and injury-associated FAP subsets** (Fig. 1G). Interestingly, trajectory inference restricted to ASCs revealed that Proliferative ASCs arise from Progenitor ASCs. Along their inferred trajectory toward Pi16^+^ FAPs, Progenitor ASCs show enrichment for key functional programs, including tissue repair, ECM remodelling, cell motility, stress adaptation, and immune modulation (Supplementary Fig. 1I). In contrast, Proliferative ASCs were primarily defined by proliferation-related transcriptional signatures, which masked other functional features.

**Figure 1:**
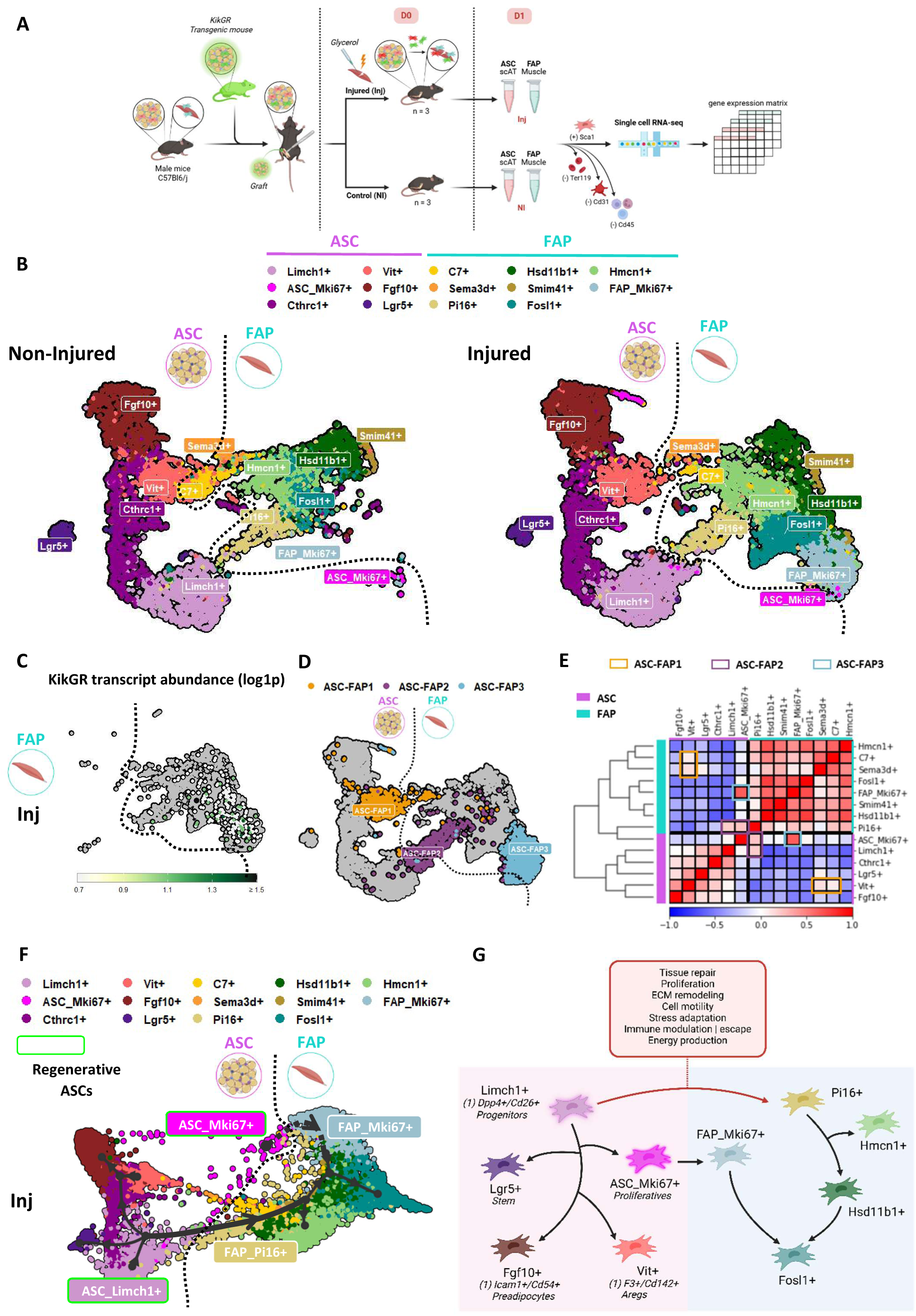
Limch1+ and Mki67+ ASC subpopulations from the scAT migrate into muscle after injury. **A** Experimental design overview of KikGR^+^ scAT grafting, muscle glycerol injury, sampling and sequencing. **B** UMAP embedding of ASCs and FAPs scRNA-seq data split by non-injured (NI) and Injured (Inj) conditions, colored by clusters. Six ASC sub-populations and eight FAP sub-populations were identified using SNN (Shared Nearest Neighbor) clustering applied to each cell type. Dotted line separate ASCs (left) from FAPs (right). **C** Feature plot showing KikGR transcript abundance in injured FAPs (FAP_Inj). **D** SNN clustering of ASCs and FAPs integration in the injured condition (Inj). Colors highlight the clusters composed of ASCs and FAPs. All other cells are colored in gray. Dotted line separate ASCs (left) from FAPs (right). **E** Correlation matrix between ASCs and FAPs sub-populations in the injured condition (Inj), organized by cell type: ASC (magenta) and FAP (blue). Colored squares highlight the clusters composed of ASCs and FAPs identified in D. **F** Trajectory inference using PAGA Tree (Partition-based graph abstraction) of the ASC and FAP sub-populations in the injured condition (Inj). Dotted line separate ASCs (left) from FAPs (right). **G** Model depicting the lineage hierarchy relationships of the ASCs and FAPs sub-populations in the injured condition (Inj). Up-regulated pathways along Limch1^+^ to Pi16^+^ trajectory are indicated.

**These observations led us to focus specifically on Limch1+ ASCs as the subset at the origin of the injury-induced migratory response, and to investigate the early transcriptional and regulatory events that govern their rapid activation in the scAT following muscle injury.**

### Injury induces a functionally primed Limch1+Prg4+ ASC subset with regenerative and adipogenic features

We next focused on the activation dynamics and early transcriptional programs of Limch1+ ASCs by performing transcriptional and regulatory analyses of ASCs from the scAT of uninjured mice (NI), at 14h (Inj_14h) and 24h (Inj_24h) post-muscle injury. We first assessed the transcriptional dynamics of ASCs following injury by comparing Inj_14h and Inj_24h to the NI state. UMAP projection revealed that ASCs exhibit an early response as soon as 14 hours post-injury, with a progressive transcriptional change of Limch1+ ASCs as well as an enrichment in Mki67+ ASCs (Fig. 2A, Supplementary Fig. 2A). These patterns and trajectories were consistent with those observed in our KikGR^+^ grafting model (Supplementary Fig. 2B). Consistent with this, differential gene expression analysis revealed a peak transcriptional response at 14 hours post-injury (Fig. 2B). Gene expression dynamics followed distinct kinetics: a subset of genes, including Mif, Cxcl14, Lgals3, Serpine2, and Ybx1, was transiently upregulated at 14h, consistent with acute inflammatory signalling, stress responses, and early tissue remodelling. In contrast, genes such as Prg4, Ssr4, Tubb5, Lgals1, Plac8, Lbp, Cxcl13, Timp1, Serpina3n, Bst2, and Fabp4 remained elevated through 24h, suggesting sustained activation of pathways linked to ECM remodelling, immune regulation, and lipid metabolism. These prolonged transcriptional programs likely support stromal remodelling and early adipogenic (*Fabp4*, *Plac8, Lgals1, Ybx1)* or fibrogenic commitment (*Timp1*, *Lgals1, Serpina3n*). **To our knowledge, this is the first evidence that ASCs initiate their activation program within the scAT prior to migration, likely facilitating their recruitment and functional engagement upon injury.** We next focused on Limch1+ ASCs to characterize their early injury response. We first performed DGE analysis between NI and Inj_14h to identify their early response to muscle injury and visualized it using a volcano plot (Fig. 2C). Limch1+ ASCs showed a stronger transcriptional response than the global ASC population, with 142 DEGs compared to 113, and higher fold changes overall. Prg4 emerged as the top upregulated gene following injury, with selective and sustained expression in Limch1+ ASCs. Beyond its canonical role in cartilage, Prg4 modulates inflammation, reduces fibrosis, promotes vascularization, and supports mesenchymal progenitor function and adipose tissue preservation during wound healing^27–29^. Notably, recombinant human PRG4 (rhPRG4) alone has been shown to enhance tissue regeneration and accelerate wound repair, highlighting its essential role^29^. Its induction suggests that activated Limch1+ ASCs may adopt a pro-regenerative secretory phenotype contributing to tissue repair. To further characterize their early activation, we performed Gene Ontology analysis on DEGs between Limch1+ ASCs and other ASC subpopulations after injury (Fig.2D, Table S1). Enriched terms included processes related to migration (e.g., *elastic fibre formation, regulation of cell shape and motility, response to hypoxia*) and regeneration (e.g., *angiogenesis, wound healing, cell activation, tissue morphogenesis*). Early induction of biological pathways such as mTOR, vitamin C/ascorbate metabolism and fatty acid were also revealed by reactome analysis (Supplementary Fig. 2D). **Together, these results suggest that Progenitor ASCs undergo rapid transcriptional and metabolic priming prior to migration.** To complement transcriptional analyses, we applied SCENIC to reconstruct gene regulatory networks (GRNs) in Progenitor ASCs. Regulons were ranked by ΔRSS, reflecting shifts in their specificity between NI and Inj_14h (Fig. 2E). Positive ΔRSS values indicated increased specificity in the injured state, revealing regulons selectively mobilized upon injury, including Prrx1 and Prrx2, known regulators of muscle regeneration. Notably, we also identified adipogenesis-associated regulons such as Ybx1, Pparg and Hmga, despite the undifferentiated state of the cells. GO analysis of target genes highlighted that injury-specific regulons were enriched in proliferation, migration, and remodelling pathways, while NI-specific ones were associated with morphogenesis and patterning. **These findings suggest that injury triggers a functional rewiring of Limch1+ ASC regulatory networks toward activation-related programs.** We further examined regulon activity by assessing AUC score distributions across Limch1+ ASCs (Supplementary Fig. 2H). Among regulons with high ΔRSS, some, such as Ybx1 and Prrx1/2, showed increased activity following injury, while others, including Pparg and Hmga2, maintained similar levels across conditions. This indicates that injury leads to the engagement of these transcriptional programs in a larger fraction of cells, with or without a change in activity intensity, thereby increasing their specificity. Transcription factors from the C/EBP family, Cebpa, Cebpb and Cebpd, were also identified as key regulators likely driving the gene upregulation observed in Progenitor ASCs at Inj_14h. These factors are well known for their role in adipogenesis, supporting the early engagement of these cells in an adipogenic program (Supplementary Fig. 2J).

**Figure 2:**
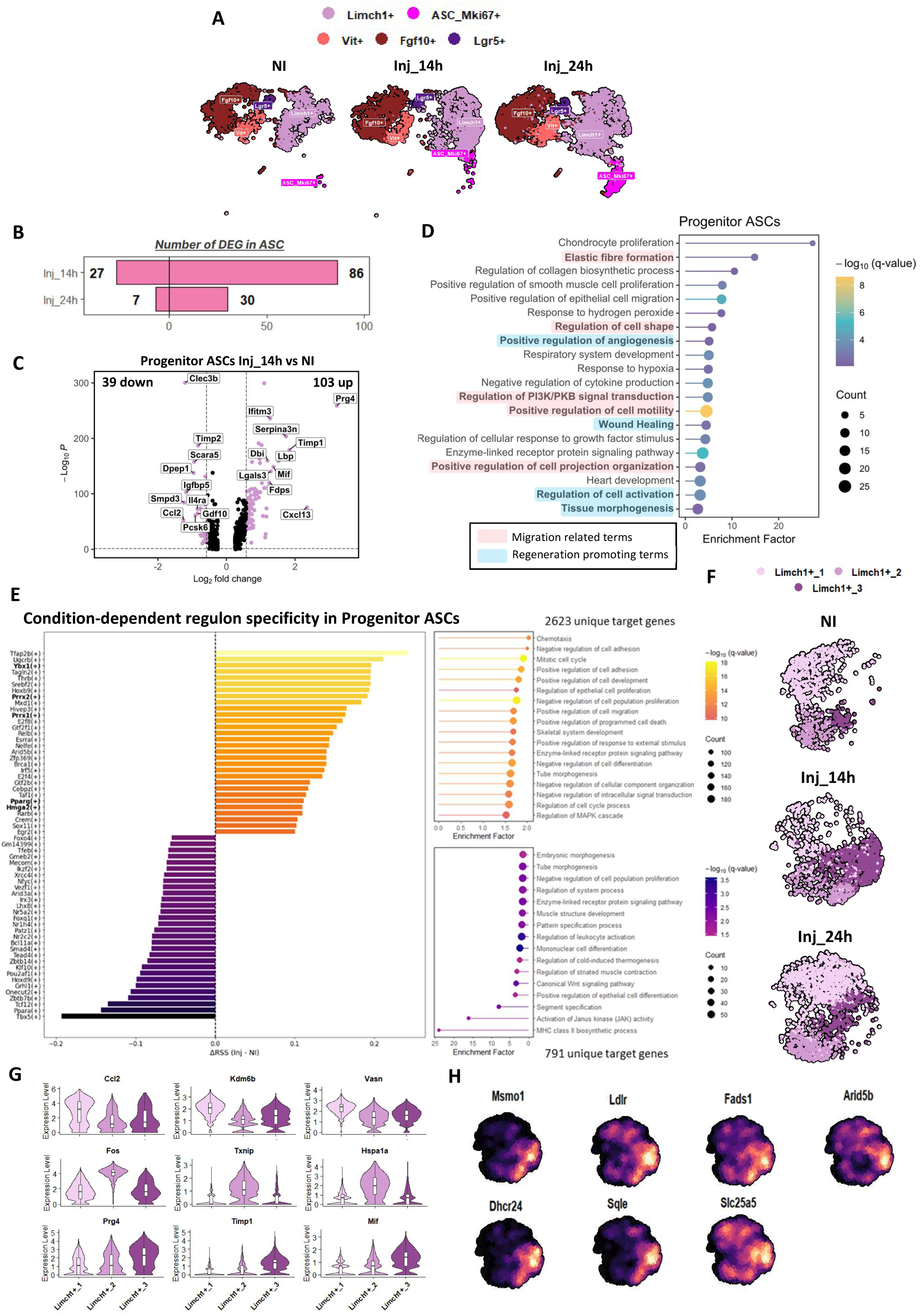
Early activation of Mobilizable-ASCs with regenerative and adipogenic potential. **A** UMAP embedding of ASCs split by non-injured (NI), 14h (Inj_14h) and 24h (Inj_24h) post-injury conditions, colored by clusters. **B** Barplot showing the number of differentially expressed genes (DEGs) in ASCs at 14 and 24h post-injury (Inj_14h and Inj_24h), compared to the nin-injured condition (NI). **C** Volcano plot of DEGs in Progenitor ASCs at 14h post-injury (Inj_14h) versus the non-injured condition (NI). Thresholds were set at log₂ fold change (log₂FC) > 0.58 and p-value < 0.05. Red points indicate genes meeting both thresholds, while blue points represent genes passing only the p-value threshold. **D** Pathway enrichment analysis of up-regulated genes in Progenitor ASCs compared to other ASC sub-populations at 14h post-injury (Inj_14h). Gene Ontology (GO), KEGG, and Reactome databases were queried. Enriched terms related to migration are highlighted in red, and those related to regeneration are highlighted in blue. **E** Condition-dependent regulon specificity in ASC progenitors. (Left) Bar plot showing ΔRSS (Inj_14h – NI) values for selected regulons inferred by SCENIC. Positive ΔRSS indicates increased specificity in the Inj_14h condition. (Right) Pathway enrichment analysis of target genes from the most condition-specific regulons, with increased specificity at the top and reduced specificity at the bottom. **F** UMAP embedding of ASC Progenitors split by conditions (non-injured –NI- and 14 or 24h post-injury -Inj_14h and Inj_24h-), colored by subclusters. **G** Violin plots showing the expression levels of representative marker genes of each Progenitor ASCs subclusters. **H** Nebulosa plots showing the expression of selected sterol metabolism-associated genes specifically upregulated in ASC Progenitors at 14h post-injury (Inj_14h). Expression density is overlaid on the UMAP of Progenitor ASCs subclusters.

**Together, differential expression and GRN analyses reveal a rapid regulatory shift in Lmich1+ ASCs, linked to regeneration, migration, and early adipogenic fate priming. Notably, the heterogeneity in regulon activation suggests the emergence of a transcriptionally distinct subset of early responders within this population.**

Given the strong injury-induced activation observed in Progenitor ASCs, we next investigated whether this population harboured early-response heterogeneity. Subclustering revealed three subpopulations (Limch1+_1–3), with Limch1+_3 emerging transiently at 14 h post-injury (Fig. 2F, Supplementary Fig. 2M). This subset expressed key injury-induced genes, notably Prg4, the top hit with potent pro-regenerative potential, alongside Timp1 and Mif, which were among the most strongly upregulated following injury and associated with inflammation, matrix remodelling, and tissue repair (Fig. 2G). Integration with FAPs across timepoints further supported this identity, revealing that Lmich1+_3 was transcriptionally closest to Pi16^+^ FAPs and Mki67+ ASCs (Supplementary Fig. 2L), **suggesting it may represent the mob-ASCs, that are Limch1+Prg4+, primed to migrate and give rise to early FAP subpopulations in injured muscle.** Cross-referencing differential expression results across all ASC subpopulations between non-injured and 14h post-injury conditions identified 31 genes specifically upregulated in Progenitor ASCs. Gene ontology analysis of these genes revealed enrichment in pathways related to cellular stress (*Chaperone cofactor-dependent protein refolding, Maintenance of protein location in cell*), migration (*Fibroblast migration, Regulation of collagen biosynthetic process*), homeostasis (*Cell redox homeostasis*) and sterol metabolism (*Sterol biosynthetic process, Cholesterol biosynthetic process, Secondary alcohol biosynthetic process*) (Supplementary Fig. 2E). Given the transcriptional evidence of adipogenic priming in Limch1+ ASCs, we next examined the expression of sterol metabolism-associated genes across subclusters. One subset, enriched at 14 h post-injury, expressed key regulators of cholesterol biosynthesis (Msmo1, Dhcr24), lipid uptake (Ldlr), and fatty acid desaturation (Fads1), along with Arid5b, Sqle, and Slc25a5 genes involved in triglyceride metabolism, lipid droplet formation, and mitochondrial energy regulation, respectively (Fig. 2H).

**This transient population, identified as Limch1+Prg4+, represents the Mob-ASCs, emerging at the peak of ASC activation and closely related to Mki67+ ASCs and Pi16+ FAPs, displays a transcriptional profile indicative of early metabolic priming. Unexpectedly, it also exhibits increased adipogenic potential, suggesting that ASC migration could contribute to the emergence of a metabolically and fate-primed population within the regenerating muscle.**

### A transcriptional signature of Mob-ASCs within muscle traces their origin and identity across tissues

To characterize Mob-ASCs within muscle, we leveraged KikGR labeling to distinguish infiltrated ASCs from endogenous FAPs at 1 dpi. A logistic regression classifier was trained using KikGR^+^ cells as positive labels. Model performance was optimized through cross-validation, testing combinations of hyperparameters including regularization strength, class weight, sampling strategy, solver, and iteration limits. Most notably, solver type and iteration number had minimal impact on classification accuracy (Supplementary Fig. 3A-C). Given the strong class imbalance (∼4% ASC vs. 96% FAP), we applied class weighting to reduce bias toward the majority class. Model performance was primarily evaluated using recall scores for ASC and FAP populations, with a focus on ASC recall to prioritize identification of ASC-derived cells. This was critical, as KikGR-FAPs still include unlabeled infiltrated ASCs. We visualized recall metrics across hyperparameter combinations to guide model optimization (Fig. 3A). We observed that high ASC recall can come at the cost of poor FAP recall, which would compromise the model’s discriminative power. Therefore, we sought to avoid extreme parameterizations that overly favored one class. We ultimately selected two candidate models based on distinct strategies (Supplementary Fig.3D) : (1) A balanced model called “Balanced”, selected to maintain balanced recall across ASC and FAP, by maximizing the lowest recall value to avoid strong bias toward either class; (2) An ASC recall-prioritized model called “Max ASC”, which first maximized ASC recall and then selected the best among those with acceptable FAP recall (>0.7). We then applied both selected models to the test sets of FAPs to assess their ability to identify ASCs. Classification performance was evaluated on this test set using multiple metrics, including ASC and FAP recall and precision, as well as macro-averaged scores (Fig. 3B, Supplementary Fig. 3E). The “Balanced” model achieved an ASC recall of 0.73 and FAP recall of 0.83, while the “Max ASC” model prioritized ASC detection (recall = 0.82) at the cost of lower FAP recall (0.74). To reduce uncertainty, we applied a stricter probability threshold (p > 0.7) to define ASC-derived cells, yielding “Balanced_corrected” and “Max ASC_corrected” versions (Fig.3B). This refinement improved FAP recall (0.89 and 0.93, respectively) and ASC precision, enhancing specificity while reducing false positives. ASC precision remained modest, in line with the expected presence of unlabelled ASC-derived cells among KikGR-FAPs. **Together, these results demonstrate that probability-filtered logistic regression signatures can reliably identify ASC transcriptional profiles within the FAP compartment, despite biological and technical variability.** We ultimately chose the “Max ASC” model as our goal is to identify in the best way possible the ASC in the FAP, which is done with a higher ASC recall, while still having a decent FAP recall. Subsequently, we projected the ASC probability scores onto the FAP UMAP to visualize their distribution across subpopulations (Fig. 3C). **Cells with the highest ASC probability were mainly found in the Fosl1+ and Mki67+ FAP clusters, previously shown to arise mostly from Mob-ASCs, further supporting their role as injury-induced subsets shaped at least partially through ASC contribution**. We next quantified the contribution of ASC-derived cells among KikGR^+^ FAPs using varying probability thresholds from the ASC classifier (Fig. 3D, Supplementary Fig.3F-G). At p > 0.5, 29.3% of FAPs were predicted as ASC-derived, a proportion reduced to 10.5% with a more stringent threshold (p > 0.7). To identify the main drivers of ASC vs.

**Figure 3:**
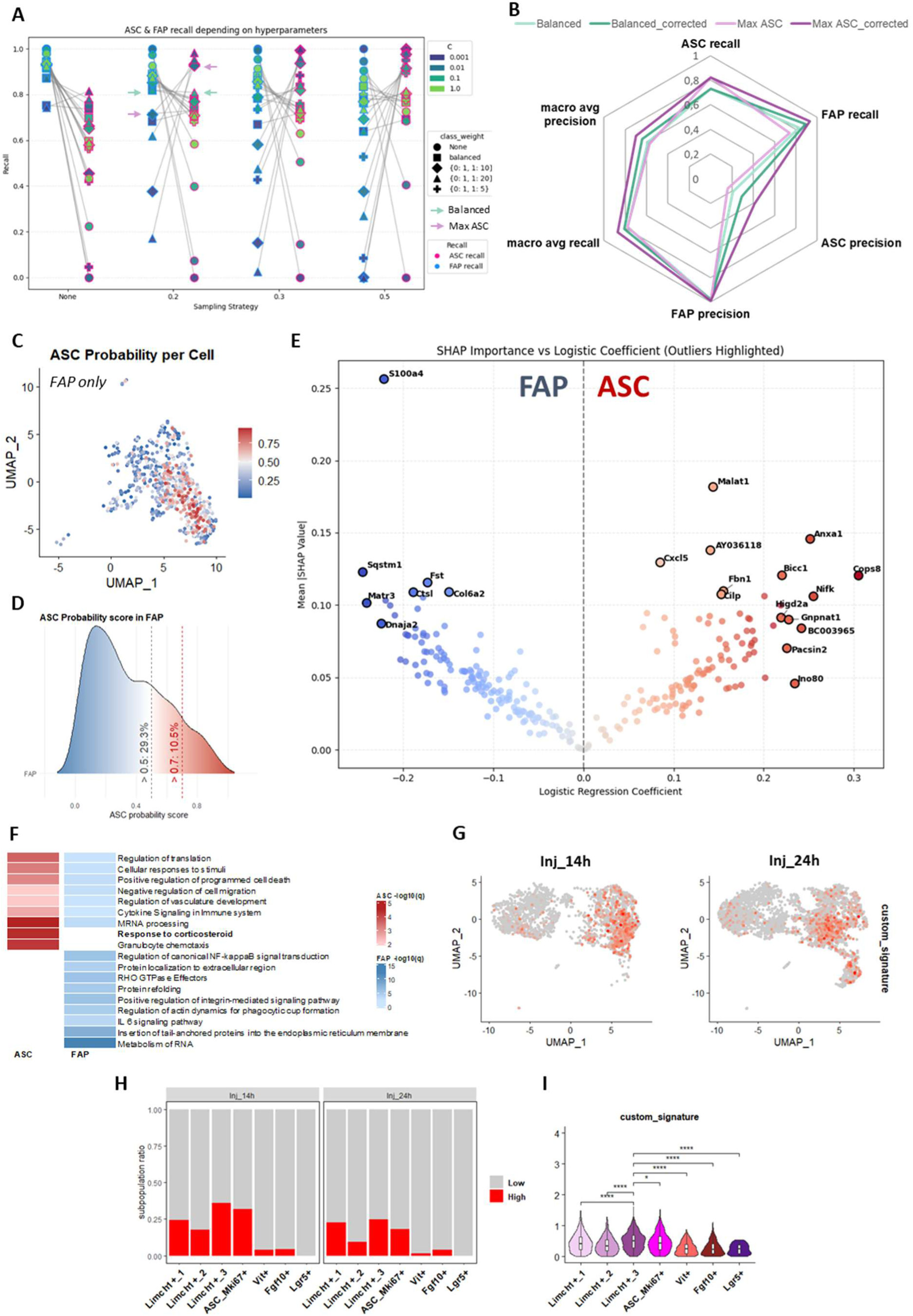
Logistic regression identifies a distinct Mobilizable-ASC signature among FAPs for tracing and functional prediction. **A** Recall scores for ASC and FAP classification across different sampling strategies and logistic regression hyperparameters. Marker shape indicates class weight, color indicates regularization strength (C), and sampling strategy is shown on the x-axis. ASC (pink) and FAP (blue) recalls are plotted separately. ASC and FAP recalls from the same set of parameters are linked with a grey line. Arrows indicate the “balanced” (green) and the “Max ASC” (purple) models ASC and FAP recall. **B** Radar plot showing classification performance on the test set using multiple metrics: ASC and FAP recall and precision, macro-averaged recall, precision, and F1-score. Curves compare original models and their corrected versions. **C** Feature plot showing ASC probability scores per cell in the FAP population in the injured condition (FAP_Inj). No KikGR^+^ cells are present in this subset. **D** Ridge plot showing the distribution of FAPs across ASC probability scores. The grey vertical line indicates the threshold of *p* > 0.5 for ASC-like classification, and the red line indicates a more stringent threshold of *p* > 0.7. **E** Volcano plot showing the relationship between mean absolute SHAP values and logistic regression coefficients for features used to predict ASC versus FAP identity. Outlier features are highlighted. **F** Pathway enrichment analysis of features used to predict ASC versus FAP identity. Gene Ontology (GO), KEGG, and Reactome databases were queried. **G** Feature plot of ASC-derived FAP signature for ASCs in the injured condition at 14 or 24h (Inj_14h and Inj_24h). **H** Bar plots showing the proportion of cells with low (grey) or high (red) ASC-derived FAP signature scores across ASC subpopulations, including Progenitor ASCs subclusters, in the injured condition at 14 or 24h (Inj_14h and Inj_24h). **I** Violin plots showing ASC-derived FAP signature scores across ASC subpopulations, including Progenitor ASCs subclusters, at 14h post-injury (Inj_14h). Statistical comparisons were performed using two-sided Mann–Whitney U tests. *p < 0.05, **p < 0.01, ***p < 0.001, ****p < 0.0001.

FAP identity, we examined both logistic regression coefficients and SHAP values (Supplementary Fig.H-J). While regression coefficients reflect the direction and strength of each gene’s association, SHAP values quantify their global contribution to classification. Plotting both revealed genes strongly associated with ASC-like or FAP-like profiles (Fig. 3E). ASC-like identity was driven by genes linked to proliferation (Malat1, Nifk, Ino80), ECM remodelling (Fbn1, Cilp, Gnpnat1), moderate inflammation (Cxcl5, Anxa family), and hypoxia response (Higd2a). In contrast, FAP-like identity was associated with genes involved in oxidative stress and autophagy (Sqstm1, Dnaja2), ECM remodeling (Ctsl, Col6a2), fibrotic activation (S100a4), and anti-inflammatory regulation (Fst). Interestingly, GO analysis of ASC-associated genes revealed selective enrichment in corticosteroid and glucocorticoid response pathways (Fig. 3F, Table S2), which are known to promote adipogenesis in both homeostatic and injury contexts^30–32^. **Altogether, these results suggest a transcriptional dichotomy between ASC-like and FAP-like identities: ASC-like cells may adopt a proliferative, remodelling-prone profile with a predisposition toward adipogenesis, whereas FAP-like cells tend to display a stress-adapted phenotype with enhanced extracellular matrix production. This contrast supports the existence of distinct functional programs between Mob-ASCs and resident FAPs during the early response to muscle injury.** To validate the identity of the Mob-ASC subpopulations identified after injury, we projected onto scAT ASC populations a score derived from the top 10 ASC-predictive and top 10 FAP-predictive genes identified by the logistic regression model (Fig. 3G). This allowed us to assess whether the transcriptional signature of Mob-ASCs is already enriched in specific scAT subpopulations prior to their recruitment. This visualization, together with the accompanying barplot showing the proportion of ASCs with high signature scores across subpopulations and timepoints, indicates that Limch1+ and Mki67+ ASCs exhibit the highest enrichment (Fig. 3H, Supplementary Fig. 3K-L). The Limch1+_3 subcluster stands out as the most enriched, with a notable decrease in signature-positive cells observed in both subpopulations by 24h post-injury. Plus, we can see that the signature score is significantly increased in Limch1+_3 but also in Mki67+ ASCs (Fig. 3I, Supplementary Fig. 3M).

**Altogether, these results confirm that Mob-ASCs Limch1+_3 ASCs but also Mki67+ ASCs are the most likely sources of mesenchymal cells within muscle, reinforcing the identity of Mob-ASCs as a transient, early activated migratory subset with regenerative potential.**

### Divergent fate commitment of FAPs during regeneration: emergence of an adipogenic fate

To explore the fate trajectories of Mob-ASCs within muscle, we investigated whether transcriptional dynamics during regeneration could reveal early lineage commitments distinguishing them from resident FAPs. Using our ASC/FAP signature, we reanalyzed single-cell RNA-seq data from Walter et al.^33^, focusing on injured FAPs from day 1 to 7 post-injury to track subpopulation dynamics and identify potential bifurcations. The clustering identified 6 different subpopulations from FAP1 to FAP6 (Fig. 4A), with an evolution in the subpopulation ratios along the regeneration kinetic (Fig. 4B). Barplot analysis revealed a marked enrichment of the FAP2 subpopulation from 3.5 dpi onward, accompanied by a reduction in FAP5, while FAP3 became dominant by 7 dpi alongside the emergence of two new subsets, FAP4 and FAP6. To assess the functional identity of these populations, we performed Gene Ontology enrichment analysis and visualized the results as a heatmap (Supplementary Fig. 4). This revealed both a shared core program and subset-specific specializations. Common pathways included *wound response, ECM organization, cell motility, vasculature development, and tissue regeneration*. FAP2 was enriched for *tissue morphogenesis* and *collagen secretion*, FAP5 for immune-related processes including *leukocyte-mediated immunity* and *IL-17 signalling*, and FAP4 for *adipocyte differentiation* and *lipid localization*. The transient emergence of adipocyte depots during regeneration, previously reported at 7 dpi^34–39^, may represent a physiological process supporting muscle repair if properly regulated^39–41^. **These findings highlight the remarkable heterogeneity of FAP subpopulations during regeneration and suggest that divergent fates are already transcriptionally specified during the initial stages of regeneration.** To validate these findings and investigate potential fate bifurcations, we applied Palantir to infer differentiation trajectories based on single-cell transcriptional dynamics. Diffusion map embedding revealed a clear separation between FAP3 and FAP4, with fate inference identifying two terminal states: fate 1 (FAP3) and fate 2 (FAP4). Cells with high branch probabilities (p > 0.7) exhibited progressive commitment toward one of the two fates, consistent with a continuous acquisition of terminal identity along each trajectory (Fig. 4D). GO analysis between the cells highly engaged in either of the two fates (Fig.4E) reveals an enrichment for *Retinol metabolism*, *Adipogenesis genes*, *Focal adhesion and TGF-β signaling* in fate 2 while fate 1 was enriched in *Regulation of actin cytoskeleton, Wnt signaling pathway, Integrin-mediated cell adhesion*. These results support fate 2 as an adipogenic trajectory, consistent with previous reports. This potential appears to be influenced by TGF-β signalling, which is known to inhibit adipogenesis. In contrast, fate 1 is associated with a non-adipogenic, likely regenerative trajectory, characterized by Wnt pathway activation, a known suppressor of adipogenic differentiation. To further validate this, we examined the expression of key adipogenic genes along pseudotime. Fate 2 showed a progressive upregulation of markers such as *Cebpd*, *Cebpb*, *Cebpa*, *Pparg*, *Klf15*, *Nr1h3*, *Lpl*, *Apoe*, and *Apod*, confirming adipogenic commitment (Fig. 4F). We also assessed *Osr1*, a marker of pro-regenerative FAPs supporting myogenic progenitors via ECM and promyogenic signals^42^. *Osr1* was predominantly enriched in fate1, aligning with renewal FAPs known to revert to a basal state after injury^7,33^ (Fig. 4G). Both trajectories showed a marked downregulation of fibrotic genes, including *Col1a1*, *Postn*, *Thbs1*, *Lox*, and *Acta2*, in line with the resolution of fibrotic activity during later stages of regeneration (Supplementary Fig.4). **These observations indicate that FAPs diverge early into distinct trajectories: one expected regenerative fate (fate 1) and one intriguing adipogenic yet regenerative path (fate 2).** To investigate whether ASCs or FAPs are predisposed toward specific fates, we projected our FAPs data with ASC and FAP labels onto the trajectories inferred from Walter et al.^33^. Remarkably, even at 1dpi, some ASCs and FAPs were already positioned along the trajectory different fates. FAPs displayed a higher proportion of cells committed to a particular fate, with a predisposition toward the classical regenerative trajectory, whereas ASCs appear to retain greater plasticity with a higher percentage of cells at the beginning of trajectory (Fig.4H, Supplementary Fig.4). Odds ratio analysis revealed that predicted Mob-ASCs within muscle were 1.36 times more likely to populate FAP4 than FAP3 compared to resident FAPs, indicating a 36% increased likelihood for ASCs to engage in an adipogenic trajectory rather than a classical regenerative fate at 1 dpi (Fig. 4I). This finding supports previous results showing their distinct responsiveness to glucocorticoids and the adipogenic priming of migrating ASCs that already occurs in the scAT. Analysis of ASC distribution along the two differentiation trajectories revealed that ASCs associated with FAP4 (fate 2) were significantly more advanced along pseudotime than those linked to FAP3 (fate 1) (mean pseudotime: 0.7 vs. 0.2; *p* < 0.0001), indicating an early and committed engagement toward adipogenic differentiation. In contrast, ASCs associated with fate 1 displayed lower fate probabilities (often < 0.6), suggesting they remain in a more plastic state and may retain the capacity to switch fate (Fig. 4J).

**Figure 4:**
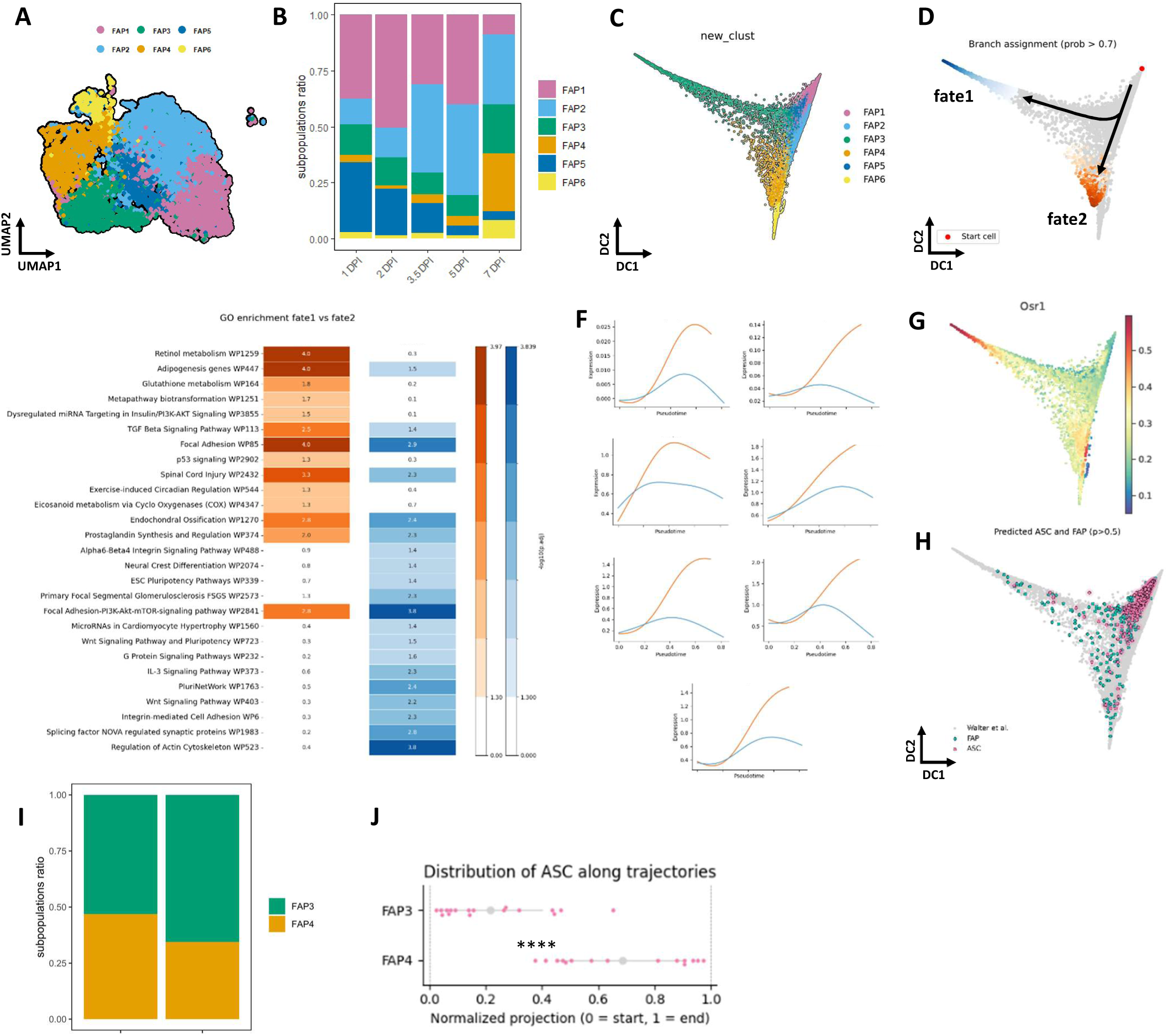
Mobilizable ASCs among FAP exhibit an adipogenic bias along divergent regenerative trajectories. **A** UMAP embedding of Walter et al. FAPs from muscle injury kinetic (1 to 7 dpi) scRNA-seq data, colored by clusters. Six FAP clusters were identified using SNN (Shared Nearest Neighbor) clustering. **B** Barplots of clusters composition of FAPs across muscle injury. **C-D** Diffusion map embedding of FAPs colored by clusters (**C**) or by predicted cell fate probabilities (**D**) using Palantir, with cells above the threshold (p > 0.7) highlighted. **E** Pathway enrichment analysis of genes differentially expressed between cells highly committed to fate 2 (orange) and fate 1 (blue), as predicted by Palantir. Enrichment was performed using the WikiPathways database. **F** Expression dynamics of key adipogenesis-related genes along pseudotime for both fate1 (blue) and fate2 (orange) trajectories. Genes were selected based on their known roles in adipogenic commitment. **G** Feature plot of Osr1 in FAPs. **H** Projection of our FAP dataset onto the differentiation trajectories inferred from Walter et al.. Cells are colored by predicted identity (ASC or FAP; *p* > 0.5). Grey cells represent the original reference dataset from Walter et al. **I** Bar plots showing the composition in FAP3 and FAP4 clusters, associated with fate 1 and fate 2, split by ASC-predicted FAPs and endogenous FAPs. **J** Scatter plot of the normalized distribution of ASC-predicted FAPs along the two differentiation trajectories represented by FAP3 and FAP4 (test de Mann-Whitney U, ****p < 0.0001).

**Altogether, these findings reveal that divergent fate trajectories within the FAP compartment emerge early during regeneration. While resident FAPs display a stronger early commitment toward a classical regenerative fate, a predisposition toward adipogenesis is also evident. In contrast, predicted Mob-ASCs within muscle, although initially less lineage-committed, exhibit early signs of engagement toward adipogenesis.**

### Intramuscular regenerative adipogenesis is transient and partially supported by infiltrated Mob-ASCs

It has been observed, although not finely characterized, that lipid deposition in the form of ectopic adipocytes occurs during the normal process of muscle regeneration^3,4,12,43^. The origin of such regenerative adipogenesis has been initially assigned to FAPs^39,44,45^ but our recent findings showing that scAT-originating ASC infiltrates the injured muscle^20^ now offer a new paradigm and force to reconsider the formation of this adipogenesis. Given the early adipogenic predisposition observed in Mob-ASCs within the muscle, we next aimed to determine whether the transient adipocytes appearing during regeneration originate from these infiltrating cells. Specifically, we investigated the presence, timing, and identity of intramuscular adipocytes in regenerating muscle, with a focus on tracing their origin back to scAT-derived ASCs. To this end, we used a glycerol-induced muscle injury model and collected muscles at multiple timepoints ranging from 5 to 28 days post-injury, followed by longitudinal sectioning, allowing us to monitor the dynamics of intramuscular adipocytes throughout the regenerative process (Fig. 5A). To visualize lipid accumulation during regeneration, we used BODIPY staining, which labels neutral lipids. Imaging revealed an increase in lipid droplets deposition at 7 days post-injury (dpi) compared to 0 and 14 dpi (Fig. 5B). Quantification of BODIPY-positive areas in longitudinal muscle sections confirmed a transient peak in lipid accumulation between 7 and 9 dpi (Fig. 5C). This observation was validated using Oil Red O staining, another neutral lipid marker, which also showed a significant increase in lipid volume, peaking at 7 dpi (Fig. 5D–E). To confirm the presence of proper adipocytes, we performed immunostaining for Perilipin, a protein coating mature lipid droplets in adipocytes. Perilipin/WGA/DAPI-positive adipocytes were clearly detectable at 7 dpi (Fig. 5F), indicating that at least part of the observed lipid accumulation corresponds to adipocytes. To assess whether ASCs contribute to this post-injury intramuscular adipogenesis, we first directly injected ASCs isolated from KikGR fluorescent transgenic mice into wild-type mouse 1h after muscle injury and analysed the muscle at 5 dpi (Fig. 5G). Imaging revealed KikGR^+^ cells filled with LipidTOX stained lipid droplets, indicating that injected ASCs were able to differentiate into mature adipocytes within the regenerating muscle (Fig. 5H). **These results indicate that ASCs brought to the injury site can differentiate into mature adipocytes.** To further determine whether this differentiation also occurs from endogenous ASCs migrating from the scAT, we grafted scAT from a fluorescent KikGR mouse into the scAT of a WT non fluorescent mouse 7 days prior to muscle injury and collected the injured muscle at 7 dpi (Fig. 5I). Imaging revealed KikGR^+^ mature adipocytes co-labelled with LipidTOX, confirming that endogenous migrating ASCs can indeed infiltrate the injured muscle and then differentiate into intramuscular adipocytes during its regeneration (Fig. 5J).

**Figure 5:**
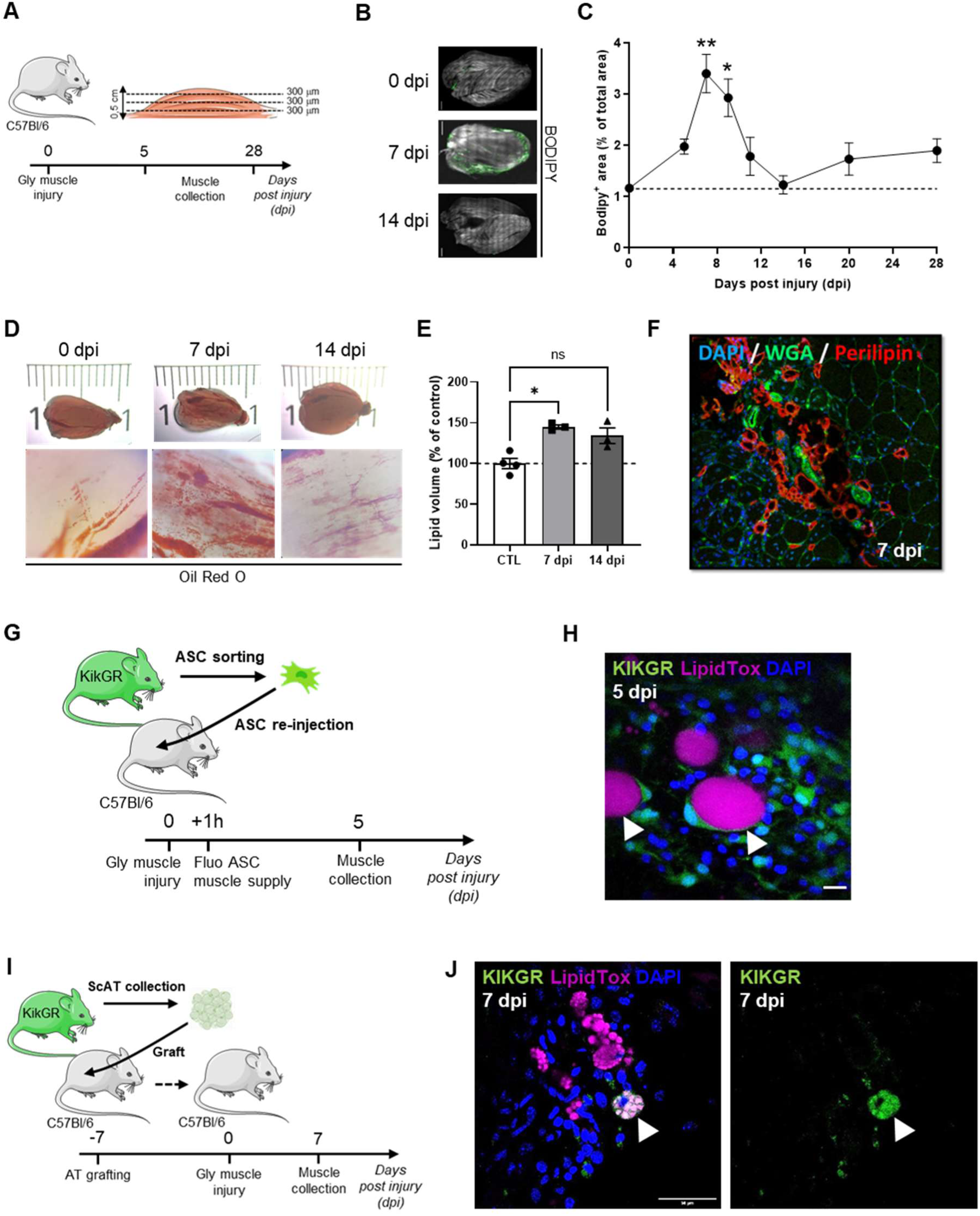
Intramuscular regenerative adipogenesis is transient and partially supported by infiltrated ASCs. **A** Experimental scheme of muscle glycerol (Gly) injury and sampling. **B** Transversal quadriceps muscle sections stained with Bodipy at 0-, 7- and 14-days post-injury (dpi). **C** Time course quantification of Bodipy positive area in muscle sections after muscle injury, each point represents the sum of 3 separated 300 μm sections of 4 to 7 animals. **D** macroscopic view of Gly-injured quadriceps transparized and stained with Oil Red O and corresponding zoom-in images at 0, 7 and 14 dpi. **E** Oil red O quantification in Gly-injured quadriceps (0, 7 and 14 dpi) of 3 to 4 animals per time point. **F** Immunohistological analysis of Gly-injured (7 dpi) quadriceps (transversal section) showing intramuscular adipocytes (WGA^+^/Perilipin1^+^/Dapi^+^). **G** Model of ASC intramuscular injection from KikGR mouse into WT C57Bl/6 mice 1h after Gly-muscle injury. **H** Immunohistological analysis of Gly-injured (5 dpi) quadriceps (transversal section) showing intramuscular fluorescent adipocytes (KikGR^+^/LipidTox^+^/Dapi^+^). **I** Model of ScAT grafting from KikGR mouse into WT C57Bl/6 mice. **J** Immunohistological analysis of Gly-injured (7 dpi) quadriceps (transversal section) showing intramuscular fluorescent adipocytes (KikGR^+^/LipidTox^+^/Dapi^+^). Bar scale 10 μm. Results are expressed as mean ± SEM; \**p* < 0.05, \*\**p* < 0.01. The figure was partly generated using Servier Medical Art, provided by Servier, licensed under a Creative Commons Attribution 3.0 unported license.

**Altogether, these results reveal a transient peak of intramuscular adipocytes between 7 and 9 dpi, that has already been observed in previous studies**^34,36,38,39,46^. Nevertheless, we demonstrate for the first time that infiltrated ASCs originating from scAT party support this transient adipogenesis. These *in viv*o observations support our *in silico* previous trajectory inference findings, which indicated that by 1 dpi, some ASCs were already transcriptionally predisposed to an adipogenic fate.

### ASC-derived adipocytes promote myogenesis

Having established that a subset of intramuscular adipocytes originates from infiltrated ASCs, we next sought to determine whether these adipocytes play a distinct role during muscle regeneration, particularly in supporting myogenesis. To investigate this, we isolated ASCs from the scAT and FAPs (their muscle counterparts) from the quadriceps muscle and cultured them *in vitro*. Adipogenic differentiation was induced in a subset of cultures to generate ASC- and FAP-derived adipocytes, named respectively Adipo_ASC_ and Adipo_FAP_. Conditioned medias (CM) were then collected from ASCs, FAPs, and their respective adipocytes, and applied to MuSCs to observe their effect on their differentiation (Fig. 6A). Immunohistological analysis was conducted after 4 days of MuSCs differentiation in the presence of various CM, using Desmin staining to identify cells of the myogenic lineage (Fig. 6B). Visually, an increase in myotube formation was observed in the condition treated with adipocyte-derived CM. To quantify this effect, we assessed multiple parameters. The fusion index, representing the percentage of myotubes containing more than one nucleus, was significantly increased across all CM-treated conditions, indicating an overall enhancement of muscle cell differentiation. Among the tested conditions, the AdipoASC one had the most pronounced effect, significantly increasing the fusion index to nearly 80%, compared to approximately 50% in the control condition (Fig. 6C). The myotube surface coverage was also markedly higher in cultures treated with adipocyte-derived CM, and particularly those from Adipo_ASC_ (Fig. 6D). The percentage of Desmin-negative cells, representing undifferentiated muscle cells, was significantly reduced in the Adipo_ASC_ condition, further corroborating the enhanced differentiation observed earlier (Fig.6E). **Together, these data indicate that the CM from Adipo_ASC_ is the one that promotes the most MuSCs differentiation**. To assess whether this differentiation was accompanied by greater myotube maturation, we analyzed the number of nuclei per myotube. Strikingly, the Adipo_ASC_ condition resulted in a significantly higher proportion of myotubes with a large number of nuclei, indicating enhanced fusion events and a more advanced maturation state (Fig. 6F-G).

**Figure 6:**
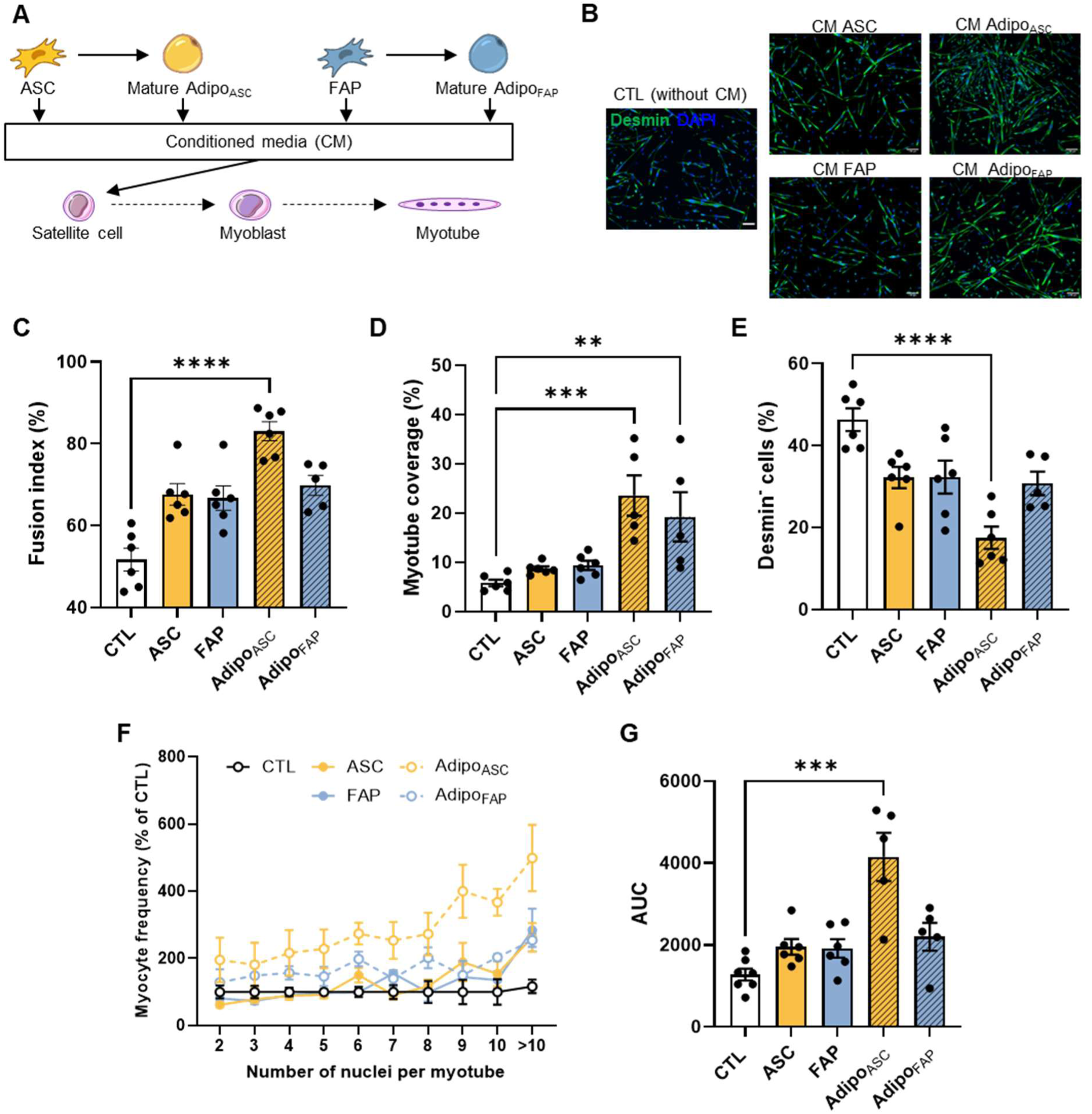
Adipocytes derived from ASCs are pro-myogenic. **A** Experimental scheme of conditioned media collection and muscle stem cell (MuSC) culture; Adipo_ASC_ and Adipo_FAP_ correspond to differentiated adipocytes derived respectively from ASC or FAP. **B** Immunohistological analysis of MuSC differentiation after 4 days of culture with the different conditioned media and stained with Desmin and DAPI, representative images. **C-E** Quantification of the myotube fusion index (C), myotube coverage area (D) and percentage of Desmin^-^ non differentiated MuSC (E) determined with MyoFInDer designed by Weisrock A et al. (doi: 10.1089/ten.TEA.2024.0049). **F-G** Myotubes repartition according to nuclei number determined with MyoFInDer. Results are expressed as mean ± SEM of 3 independant experiments; \*\**p* < 0.01, \*\*\**p* < 0.001, \*\*\**p* < 0.001 and \*\*\*\**p* < 0.0001.

**These findings demonstrate that among all tested populations, including undifferentiated ASCs, FAPs, and Adipo_FAP_, Adipo_ASC_ secretion emerged as the most potent in promoting MuSCs fusion and myotube maturation, highlighting their distinct pro-myogenic capacity.**

## Discussion

Muscle regeneration is accompanied by a rapid activation of resident mesenchymal progenitors called FAPs, with a proliferation peak typically observed around 3dpi. Intriguingly, several studies have reported a fivefold increase in FAP numbers within the first 24 hours after injury, prior to the onset of proliferation. Recently we demonstrated that this early expansion is due to the recruitment of ASCs derived specifically from the scAT, which migrate into the injured muscle and contribute to the FAP compartment^20^. Despite this discovery, the identity of these migrating ASCs, the mechanisms governing their activation, and their specific role within the regenerating muscle remain unknown.

Our current study demonstrates that among ASCs, a specific subpopulation, the Limch1+ ASCs, is rapidly and selectively activated following injury to give rise to Limch1+ Prg4+ ASCs that we call the Mobilizable ASCs (Mob-ASCs) that then migrate into the muscle where they contribute to the emergence of injury-associated FAP subpopulations. Notably, muscle injury induces a transient expansion of a functionally primed subset within this population, characterized by both regenerative and adipogenic features. This subset likely corresponds to the specific Mob-ASCs recruited upon injury, carrying an intrinsic predisposition toward adipogenesis. By tracking marked ASCs within the FAP compartment, we predicted their transcriptional signature, enabling the identification of Mob-ASCs among resident FAPs. This signature revealed enhanced sensitivity to glucocorticoid signaling, a pathway known to promote adipogenic commitment^30,31,47^. This suggests that ASCs primed in the scAT retain a tendency to differentiate into adipocytes after entering the muscle, likely reinforced by local glucocorticoid signals. At 1 dpi, Mob-ASCs are more plastic than resident FAPs and more prone to engage in adipogenesis.

Interestingly, FAPs contribution to the transient formation of ectopic adipocytes during muscle regeneration have already been described^3,4,12,43^. This mechanism has long been seen as detrimental because involved in intramuscular adipose tissue (IMAT) development in pathologies^45,48–50^. However, under normal conditions, it remains a controlled and transient phenomenon that may represent a physiological process of muscle regeneration^39–41^, as genetic ablation of committed or differentiating muscle adipocytes leads to impaired regeneration, characterized by persistent inflammation, reduced MuSCs expansion and number and size of regenerated myofibers^40,51^.

Using GLY-induced injury, which leads to greater IMAT formation than CTX-induced injury^39^, we show for the first time that Mob-ASCs from scATcontribute to the transient adipocyte peak during regeneration. Among all tested populations, adipocytes originating from ASCs most effectively promoted myogenic cell maturation, surpassing the effects of both FAP-derived adipocytes and undifferentiated ASCs or FAPs, suggesting that ASC migration supports muscle regeneration not only by replenishing the FAP pool but also by triggering a transient, pro-myogenic adipogenic response.

Our findings align with previous studies showing that intramuscular adipocytes support muscle regeneration, and that a transient adipogenic response is a hallmark of successful recovery. Notably, Kuang et al. demonstrated that genetic ablation of adipogenic progenitors impaired myofiber regeneration in CTX-injured muscles, despite the lower IMAT levels induced by CTX compared to glycerol, underscoring the importance of even modest, transient adipogenesis in effective muscle repair^51^.

Similarly, Feige et al. showed that loss of the adipogenic regulator PPARγ in a GLY-induced injury model on tibialis anterior (TA) completely abrogates ectopic adipogenesis during regeneration and leads to defective expansion and function of MuSCs^40^. Together, these studies support the concept that transient adipogenic responses and IMAT formation are critical components of the regenerative cascade.

Nevertheless, excessive or persistent IMAT can compromise muscle function. Using a GLY-induced injury model on the extensor digitorum longus (EDL) muscle, Meyer et al. demonstrated a direct negative impact of IMAT accumulation on muscle contractile properties at 21 dpi, independent of changes in contractile cross-sectional area^50^. They later showed that the complete absence of AT severely impairs muscle homeostasis. In fat-free mice, muscle mass and contractile function were restored by reconstituting only ∼10% of normal adipose mass, an effect entirely dependent on adipose-derived leptin^52^. Although not performed in an injury context, this clearly demonstrates that adipocyte-derived signals are essential for normal muscle physiology.

A recent study by Norris et al. used a mouse model in which *Pparγ* is deleted specifically in Pdgfra^+^ cells, to prevent FAP-driven IMAT formation after an adipogenic injury^44^. They reported that blocking IMAT formation enhances regeneration, but the IMAT levels in their model are markedly higher and non-transient compared to other studies^39^, including ours. This suggests their findings reflect suppression of pathological IMAT, rather than evidence that physiological IMAT is detrimental. Even more, residual IMAT in their model remains within normal levels and does not impair repair, supporting the idea that moderate, transient IMAT may be compatible with efficient regeneration.

It is why it is important to consider that IMAT formation, composition, and role may vary depending on factors such as sex, strain, age, muscle type, and type of injury^46^. Baseline FAP percentages, proliferation capacity, and adipogenic potential can differ significantly, indicating intrinsic differences among FAPs from distinct anatomical regions^53^. Another point to take into account is that adipocytes are heterogeneous^54–57^, and their composition may also vary depending on the muscle type and their origin. In this study we demonstrated for the first time that the transient peak of adipocytes observed during muscle regeneration includes ASCs derived from scAT, which display higher pro-myogenic properties. We can therefore hypothesize that depending on their heterogeneity, IMAT may be more or less deleterious or beneficial for muscle regeneration. Performing bulk RNA-seq or snRNA-seq on ASC- and FAP-derived adipocytes, would allow us to assess their endogenous heterogeneity and determine whether these ectopic adipocytes belong to known subpopulations^54–57^ or represent a novel pro-regenerative adipocyte type. Interestingly, a study in human comparing adipocytes derived either from ASCs prepared from the scAT or from FAPs from muscle shows that FAPs give rise to white adipocytes that have the unexpected feature of being insulin-resistant compared to ASC-derived adipocytes^58^. Which is interesting as adipokines secretion, known for their role in myofiber maturation, are stimulated by insulin^59,60^. It may therefore be relevant to investigate if insulin-resistant adipocytes, derived from local FAPs, might exhibit reduced secretion of lipo-cytokines, thereby attenuating their potential beneficial effects during regeneration compared to ASC-derived adipocytes.

Interestingly, in our study, ASC-derived adipocytes appeared more pro-myogenic than FAP-derived ones, also suggesting intrinsic differences that merit further investigation. Based on our initial hypotheses, these adipocytes may support regeneration by providing metabolic fuel for differentiating MuSCs, lipid precursors such as phosphatidylcholine and serine for membrane synthesis, and pro-myogenic adipokines, a possibility supported by our observations. Investigating ASC- and FAP-derived adipocytes heterogeneity, lipidomic profile, and secretome could provide insight into these hypotheses.

In summary, our findings establish that the transient and controlled appearance of adipocytes during muscle regeneration, partly composed of ASC-derived adipocytes, is not a pathological event but rather a physiological and pro-regenerative process. We show that Limch1+Prg4+ ASCs, the Mob-ASCs, rapidly responds to injury by migrating into the muscle, where it gives rise to injury-associated FAPs predisposed to adipogenic differentiation. This transient adipogenic peak, potentially reinforced by local glucocorticoid cues, supports muscle repair, with ASC-derived adipocytes being particularly pro-myogenic and promoting myofiber maturation. This view is consistent with previous studies showing that genetic ablation of differentiating adipocytes impairs regeneration, but we further provide a more detailed characterization of adipocyte kinetics, their origin, and specificity. It also challenges the idea that IMAT is intrinsically detrimental, as proposed in models where adipogenesis leads to unphysiological IMAT levels without the transient peak typically observed in successfully regenerating muscle, which instead resembles a “chronic-like” adipogenesis. Together with the literature, our data highlight that IMAT acts as a context-dependent regulator: beneficial when its formation is transient and quantitatively controlled, but deleterious over a certain threshold, not defined yet, when it becomes excessive or persistent. Importantly, we demonstrate for the first time that ASC-derived adipocytes display a stronger pro-myogenic potential than those derived from FAPs or undifferentiated ASCs and FAPs, making them a promising target to harness for improving muscle regeneration in pathological settings.

## Methods

### Animal experimental protocols

This work was submitted to and approved by the Regional Ethic Committee and registered to the French Ministère de la Recherche. Animals were kept under controlled light (12-hr light/dark cycles; 07h00–19h00), temperature (20 °C–22 °C), hygrometry (40% ± 20%) and fed ad libitum a chow diet (8.4% fat, Safe®A04, Safe lab).

### Muscle injuries

8-12 weeks old male C57Bl/6 J mice (Janvier) were anesthetized with isoflurane and 80 µL 50% glycerol in saline solution (NaCl 0.9%) were injected into the right quadriceps.

### ASC intramuscular injection and scAT grafting

The transgenic fluorescent mouse model used as ASC or scAT donor is the CAG::KikGR model referred as KikGR mice. Non fluorescent C57Bl/6 J-recipient mice received either intramuscular injection of sorted fluorescent ASC isolated from KikGR scAT (250,000 to 300,000 freshly isolated ASCs injected with G25 needle, 1 mL syringe in NaCl 0.9%, maximal volume of 80 µl in anesthetized animals 1 hour after muscle injury), or scAT graft pads (10 mg) from the above model into their scAT while sham control mice were only skin incised. After 7 days (grafted or sham) mice were injured.

### Tissue collection

Mice were euthanized via dislocation. ScAT were either directly harvested for cell isolation or fresh frozen for RNAs extraction. Quadriceps muscles were directly harvested for cell isolation, fresh frozen for RNAs extraction or fixed for further histological studies (see other sections).

### Murine AT- or muscle-SVF isolation

Freshly harvested AT or muscle were minced and SVF were obtained by enzymatic digestion as described in Sastourné-Arrey et al.^20^

### Cell sorting

ASCs were isolated from scAT and sorted with magnetic beads as described in Sastourné-Arrey et al.^20^. Isolated ASCs were used for RNAseq experiments, cell culture and conditionned media collection, muscle injection.

Muscle stem cells (MuSCs) were isolated from quadriceps muscle. Derived SVF were depleted in CD45+ and CD31+ cells using anti-CD45-FITC (Miltenyi, 130-116-535) and anti-CD31-FITC antibodies (Miltenyi, 130-123-675) followed by anti-FITC magnetic microbeads (Miltenyi, 130-048-701) using an autoMACS® Pro Separator (MACS Cell Separation, Miltenyi Biotec SAS) according to the manufacturer’s instructions. CD45^−^/CD31^−^ cells were then positively sorted for Sca-1 with anti-Sca-1 magnetic microbeads (Miltenyi, 130-106-641). CD45^−^/CD31^−^/Sca-1^-^ cells were finally positively sorted for α7-integrin with anti-α7-integrin magnetic microbeads (Miltenyi, 130-104-261). Isolated MuSCs were used for cell culture experiments.

### Cell culture

ASCs and FAPs were cultured in multi-well dishes at 37°C in a 5% CO2 and 5% O2 atmosphere. They were seesed at a density of 60000 cells/cm^2^ in αMEM (90%), NCS (10%). Media is changed every other day.

For adipogenic induction, ASCs and FAPs were brought to ∼80% of confluence, and were then cultured in pro-adipogenic media: αMEM (98%), NCS (2%), Dexamethasone 33nM, Insulin 5 mg/ml), Rosiglitazone (1 μM), T3 (10 μM), Transferrin (10 μg/ml). After 4 days of incubation, medium as changed.

Satellite cells (SC) were cultured in multi-well dishes coated with 0.3% rat tail collagen (Corning #354236) at 37°C in a 5% CO2 and 5% O2 atmosphere. SC were seeded at a density of 20000 cells/cm^2^ in DMEM / Glutamax 1g/L glucose (40%), HAMF10 (40%), Fetal bovine serum (20%) + 12.5ul bFGF (2.5ng/mL). After 7 days, when ∼70–80% of confluence was reached, cells are trypsinised and reseeded in multi-well dishes coated with Matrigel at a density of 50000 cells/cm^2^ in DMEM / Glutamax 1g/L glucose (49%), HAMF10 (49%), Fetal bovine serum (2%) to start myogenic differentiation. If needed, media is changed every other day.

For SC proliferation assay, cells seeded at 10000 cells/cm^2^ in 96-well plate and were cultured as mentionned previously for 48h in the Incucyte S3® Live-Cell Analysis System (Sartorius) enabling real-time, automated cell proliferation assays.

### Conditioned media experiments

For collection of ASC and FAP derived conditionned media, cells were first brought to ∼80% of confluence as described eralier. Then, ASCs and FAPs were rinced with PBS and cultured for 24h in a neutral medium HAMF10 (50%), DMEM glutamax 1g/L glucose (50%). Medias were collected, centrifugated to remove cell debris and strored at -80°C until use. ASC and FAPs are cultured back with proliferation media for another 24h. Pro-adipogenic media is then added and cells were differentiated in to mature adipocytes for 5 days. Identically, ASCs and FAPs-derived adipocytes were then cultured another 24h in neutral medium before media collection. As a result, we collected paired-secretions of the same cells before and after their adipogenic differentiation.

### Immunofluorescent Staining

Cells were cultured in multi well dishes and fixed with 4% paraformaldehyde. Samples were incubated at RT in permeabilization medium (0.1% TritonX-100 in Phosphate Buffer Solution, PBS), for 10 min, followed by blocking with 0.1% new horse serum, 0.1% Tween20 in PBS, at RT for 1h. Cells were incubated overnight with Desmin primary antibody (Cell Signaling) 1:100 in 0.1% new horse serum, 0.1% Tween20 in PBS at 4°C overnight. Before applying the appropriate secondary antibody (1:500, Alexa Fluor® donkey 488 anti-rabbit, Thermo Fisher Scientific), the cells were washed twice with PBS. DAPI (1:3000 in PBS, Thermo Fisher Scientific) was added for 10 min and the cells were washed twice with PBS. For fusion index (ratio between myonuclei and total number of nuclei in a field), we counted the nuclei in desmin positive myotubes composed of at least 2 nuclei. This was assessed in 2 pictures (center and edge of each well) for every well in pictures taken at 10x magnification. Visualization occurred with an Eclipse Ti Microscope (Nikon) and analyzed with opensource softwares Myotube analyzer (https://github.com/SimonNoe/myotube-analyzer-app) and MyoFInder (https://tissueengineeringlab.github.io/MyoFInDer/usage.html).

### Oil Red O Staining to Assess intramuscular lipid Deposition

For intramuscular lipid deposition quantification, muscles were first transparised with SDS 1% for 72h and then fixed in PFA 4% for 36h before extensive washing in PBS. Muscles were washed twice with isopropanol 60% for 10 min and incubated with Oil Red O solution (65% of 0.5% w/v Oil Red O in isopropanol (Thermo Fisher Scientific) and MQ water) for 30 min. They were then washed extensively with SDS 1% for 8h and PBS. To quantify Oil Red O staining, lipids were extracted with isopropanol 100% for 12h and quantified for their absorbance at 510 nm for 0.1 s with Victor Spectrophotometer (PerkinElmer). Standard curve was applied and quantification expressed as a percentage of the absorbance observed in TD samples.

### Histological Analyses

For muscle fiber analysis, half quadriceps muscles were embedded in OCT before being frozen in isopentane cooled in liquid nitrogen and stored at −80°C for further analyses. 10 μm transversal muscle sections were obtained using a CryoStarTM NX70 Cryostat (Thermo Fisher Scientific), while kept at −20°C. Sections were immediately fixed in PFA 4% for 10 min. After washing with PBS, slides were permeabilized in PBS 1X, 0.2% Triton X-100 for 10min and further blocked in PBS-T, new horse serum 3% for 1h at RT. Plin1 primary antibodies (Cell Signaling, 1:250 in PBS-T) was applied overnight at 4°C. After extensive washing with PBS, muscle slides were incubated for 1 h at RT with appropriate secondary antibody (1:200 in PBS-T, Alexa Fluor® 594 donkey anti-rabbit, Thermo Fisher Scientific). WGA 488 (1:500 in PBS-T, Thermo Fisher Scientific) was added at the same time. Dapi (1:10000 in PBS, Thermo Fisher Scientific) was added for 10 min and the samples were extensively washed with PBS before mounting with ProLong® Gold antifade reagent (Molecular Probes). Visualization and whole muscle section mosaics were performed using an inverted AxioObserver videomicroscope (Zeiss) using Zen Blue software. Fiber muscle analysis (centronucleated fibers, fiber size) and Plin1 signal analysis were then performed with internal macro of ImageJ.

For intramuscular global lipid deposition and intramuscular ASC-derived adipocyte observation, entire quadriceps were fixed in PFA 4% for 24h. After washing with PBS, muscles were embedded in agarose 3% before being longitudinaly sectionned (300 μm slices). Tissues were then incubated in Bodipy 493/503 (1:500 in PBS, Thermofisher Scientific) or HCS LipidTOX^TM^ Deep Red Neutral stain (1:500 in PBS, Invitrogen) for 40 min. Dapi (1:10000 in PBS, Thermo Fisher Scientific) was added for 10 min and the samples were extensively washed with PBS before imaging with LSM880 Confocal microscope (Zeiss) using Zen Black software. For intramuscular global lipid deposition assesment, the ratio of the Bodipy positive area of the entire muscle section of 3 longitudinal sections were analysed.

### RNAs, RT-qPCR

Extraction was performed on frozen cells or whole tissues using RNA Extractions Mini kit (QiaGEN) following manufacturer’s instructions. Briefly, samples were unfrozen in RLT and lysed with tissue lyser® (QIAGEN). Samples were passed through columns with washing steps to purify RNAs. Elution was performed with RNAse free water, and RNAs concentration was evaluated with Nanodrop® 2000c (Thermo Scientific). For qPCR, RNAs were reverse transcripted using High-capacity reverse transcriptase (Invitrogen) and diluted at 50 ng/μL in RNAsefree water. Then, qPCR was performed using Fast SYBR® Green Mix (Applied Biosystems) on 96/384 wells plate, results were acquired with StepOne v2.3-3 and Viia7 devices (Life Technologies) and data were analyzed with Real-Time qPCR Studio (Life Technologies) using the 2−ΔCt method compared to Ctrl condition.

### Statistics

Data are expressed as the mean ± s.e.m. Statistical analyses were performed using Student’s t test (two-sided paired or unpaired) or oneway ANOVA followed by the post hoc Dunnett’s and Tukey’s test with GraphPad Prism software v9 (GraphPad Software). P values less than *p < 0.05; **p < 0.01 and ***p < 0.001 were statistically significant.

### KikGR amplification

A triple-nested PCR approach was used to amplify the KikGR transcript from our ASC_Inj, FAP_NI and FAP_Inj single-cell cDNA libraries with high sensitivity and specificity. The approach used KikGR-specific reverse primers combined with generic forward primers that preserves the single-cell barcoding structure. In total, there were 4 total PCR steps (3 nested steps that increasingly filtered for a specific transcript and with the last ones that added a Read2 sequence, and 1 step that added an Illumina P7 adaptor sequence). A sample from ASC without KikGR cells was used as a negative control.

qPCR was performed using Phusion Plus PCR Master Mix (Thermo Scientific) on 96 wells plate, results were acquired with StepOne v2.3-3 device (Life Technologies) and data were analyzed with Real-Time qPCR Studio (Life Technologies).

After each of the nested steps, a SPRIselect (Beckman Coulter) purification was performed using a Left Side Size Selection, whereby magnetic beads were added at a 0.8x ratio after the 1^st^ PCR and at 0.95x ratio after the 2^nd^ PCR, to bind and retain larger DNA fragments. This selection was done to enrich for DNA fragments within the desired size range and eliminate undesired products to ensure high-quality input for subsequent steps.

In Step 1, a PCR was conducted using a forward primer containing both the Illumina P5 sequence and part of the Read1 sequence (sc3’) and a reverse primer (sc2) containing a KikGR-specific sequence approximately ∼347bp upstream of the polyA tail, for a total size of 462 bp.

In Step 2, a PCR was conducted using the same forward primer that in Step 1 and a reverse primer (scN3) containing a KikGR-specific sequence with a Read2 floating tail, approximately ∼154bp upstream of the polyA tail, for a total size of 269 bp.

In Step 3, a PCR was conducted using the forward primer from Step 1 and a reverse primer (scN4) containing a KikGR-specific sequence with a template for the index primer (Read2) as a floating tail, approximately ∼91bp upstream of the polyA tail, for a total size of 206 bp.

In Step 4, a PCR was conducted using the forward primer used in Step 1 and a reverse primer containing the Read2 sequence combined with the Illumina P7 sequence to prepare cDNAs for sequencing.

Informations about primers can be found in Table S3.

Quality control of each step was verified using a Fragment Analyzer system. The different libraries were sequenced on an Illumina NextSeq 550 sequencing machine. The sequencing cycle settings were as follows: 28 cycles for Read 1, 8 cycles for the i7 index, and 91 cycles for Read 2.

To identify individual cells in the scRNA-seq library as KikGR positive, we analyzed the files of the amplicon libraries. Alignment quality of reads mapping to the KikGR sequence was evaluated by extracting alignment scores (AS tags) of KikGR from the BAM file. For example, among the 625,402 reads mapped to KikGR for FAP_Inj, AS values ranged from 42 to 88, with a median AS of 87. Approximately 94% of reads displayed an AS of 87 or 88, indicating high-quality and nearly perfect alignments. Only about 0.4% of reads had AS values below 70, suggesting minimal presence of poorly aligned sequences. Cells with at least 1 KikGR read were marked as “KikGR+”.

### Single-cell RNA sequencing and read processing

Single-cell suspensions were loaded onto a Chromium Controller (10x Genomics) using the Single Cell 3′ v3 chemistry, with a target capture of 4000 cells per condition. Single cell barcoded droplets (GEMs) were generated according to the manufacturer’s protocol. Following reverse transcription and cDNA amplification, KikGR libraries (ASC_NI, ASC_Inj, FAP_NI, and FAP_Inj) and No graft libraries (scAT_NI, ASC_Inj_14h, scAT_Inj_24h and Muscle_Inj_14h) were prepared following the 10x Genomics protocol and sequenced on an Illumina NextSeq 550 at the “Genome and Transcriptome” (GeT) platform (Genotoul, Toulouse, France).

The Cell Ranger single-cell software (version 7.0.1) was used to perform sample alignment to mm10 (GRCm38) mouse genome on the resulting reads, filtering, and UMI counting. For KikGR experiment, the KikGR gene was added to the FASTA and GTF files.

### Single-cell RNAseq analysis

Downstream analysis was performed in R (v4.2.0). Quality control, filtering, normalization, clustering, visualization, and differential expression analyses were carried out using the Seurat package (v4.1.1) with adaptations to the standard pipeline.

### Quality control and filtering

***<u>KikGR experiment:</u>*** Preprocessing was performed by celltype, ASC (NI and Inj) and FAP (NI and Inj). ASC conditions were preprocessed together given their similar distributions, while FAP were preprocessed by condition. For each dataset, an initial unsupervised analysis was conducted without filtering to assess the distribution of UMI counts, gene features, and mitochondrial content across clusters, to identify and remove low-quality cell populations or sequencing artifacts.

Cells were retained based on dataset-specific thresholds:

- ASC_NI and ASC_Inj: >4,000 UMIs, >2,000 detected genes, and <5% mitochondrial reads.
- FAP_NI: >4,050 UMIs, >1,500 detected genes, and <10% mitochondrial reads.
- FAP_Inj: >4,500 UMIs, >1,800 detected genes, and <7% mitochondrial reads.

KikGR+ cells were identified using metadata annotation prior to removing the *KikGR* gene from the count matrix to prevent downstream bias.

To identify and exclude potential doublets, we applied the *scDblFinder* R package (v1.22.0) via Bioconductor (Release 3.21) on each dataset prior filtering.

***<u>No graft experiment:</u>*** Preprocessing was performed by tissue, scAT (NI, Inj_14h and Inj_24h) and Muscle (Inj_14h). Cells were retained based on tissue-specific thresholds, which were more permissive due to celltypes diversity:

- scAT: >500 UMIs, >500 detected genes, and <15% mitochondrial reads.
- Muscle: >1300 UMIs, >500 detected genes, and <10% mitochondrial reads.

Doublets were also removed using *scDblFinder* R package (v1.22.0) via Bioconductor (Release 3.21) by tissue.

Subsetted ASC and FAP from those tissues undergo another quality check adapted to their celltype:

- ASC: >1000 detected genes and <7% mitochondrial reads.
- FAP: >1200 detected genes and <7% mitochondrial reads.

### Integration, clustering and annotation

***<u>KikGR experiment:</u>*** ASC and FAP datasets were first analysed independently. Within each cell type, conditions (NI and Inj) were normalized separately using *SCTransform* (v1.0.1) regressing out the percentage of mitochondrial UMI counts. They were then integrated with canonical correlation analysis (CCA) by condition.

Following integration, dimensionality reduction was performed using principal component analysis (PCA), and the first 40 principal components (PCs) were used for downstream clustering and visualization. Unsupervised clustering was performed using the Shared Nearest Neighbor (SNN) modularity optimization-based method implemented in Seurat with a resolution of 0.4. Cell embeddings were visualized using Uniform Manifold Approximation and Projection (UMAP) for non-linear dimensionality reduction. Markers for ASCs and FAPs were identified using a non-parametric Wilcoxon rank sum test with P values adjusted using Bonferroni correction, the markers were considered significant if they exhibited a log₂ fold-change > 0.58 and were expressed in at least 10% of the cells in the cluster, they were part of the most up-regulated genes in each clusters, they were also evaluated based on the distinctness of called markers. The clusters were annotated based on marker genes. ASC and FAP datasets were then merged and re-normalized by condition regressing out the percentage of mitochondrial UMI counts. They were then integrated with CCA by condition, re-processed using the above method and clustered with a resolution of 0.5.

***<u>No graft experiment</u>*:** scAT and Muscle datsets were analysed independently. The scAT and Muscle were normalized using *SCTransform* regressing out the percentage of mitochondrial UMI counts, integration was not needed for scAT. Dimensionality reduction was performed using PCA, and the first 40 PCs were used for downstream clustering and visualization. Unsupervised clustering was performed using the SNN modularity optimization-based method implemented in Seurat with a resolution of 0.13 for scAT and 0.2 for Muscle. Cell embeddings were visualized using UMAP for non-linear dimensionality reduction. Clusters identified as ASC and FAP, respectively for scAT and Muscle, using marker genes, were subset into individual objects, and re-processed using the above method. Clustering was performed using a resolution of 0.18 for ASC and 0.5 for FAP, to obtain similar resolution as in the KikGR experiment. Markers for ASC and FAP clusters were identified as above.

Progenitor cluster from ASC was subclustered. It was first subset then re-processed using the above method. Clustering was performed using a resolution of 0.18.

### Gene pathway analysis

Analysis of enriched pathways was performed using Metascape (v3.5.20250101). Genes selected for analysis were filtered to a Benjamini–Hochberg adjusted P-value < .05 and a log₂ fold-change > 0.58, then evaluated for enrichment in core set of ontologies for enrichment analysis, including GO processes, KEGG pathways, Wiki pathways, Reactome gene sets, canonical pathways, and CORUM complexes. Pathways and log10(adjusted p-values) can be found in Supplementary Table.

### Literature dataset analysis

Datas were obtained from Walter et al.^33^, in the GEO under accession GSE143437 (2 dpi, 5 dpi, 7 dpi) and GSE232106 (1 dpi, 3.5 dpi). Datasets from young mice were subset to contain only FAPs. They were normalized using *SCTransform* and integrated by timepoints using RPCA integration with 3000 variable genes. The UMAP reduction was calculated using the top 40 PCs and FAP clustered with a resolution of 0.25.

### Trajectory inference and RNA velocity

Trajectory inference across ASC and FAP populations was performed for injured conditions, using PAGA Tree implemented through the dynverse R toolkit^61^. The stability of the inferred trajectory was confirmed by varying model parameters (number of neighbors and resolution), which did not substantially alter the trajectory topology. The analysis was initialized with a fixed random seed (seed = 17) to ensure reproducibility. Cells were embedded using a force-directed layout, and the PAGA Tree algorithm was applied to infer a branching topology representing potential developmental paths. The root node was selected based on the ASC_Limch1+ subpopulation, and trajectory probabilities and pseudotime values were computed for each cell. Branch assignment and lineage progression were then visualized and analyzed. Gene expression dynamics along pseudotime were visualized using branch-specific heatmaps generated by the dynverse workflow. For each inferred trajectory, the top pseudotime-associated genes were selected based on expression variability and fitted with smooth trends. Genes were clustered by temporal expression pattern and visualized as heatmaps to highlight branch-specific transcriptional modules. To validate the inferred directionality of the PAGA Tree trajectory, we applied uniTVelo (v0.2.5.2) on the AnnData object after selecting the top 2000 highly variable genes. Cell-cell neighborhood graphs were computed using 30 principal components and 30 nearest neighbors. The model was trained using default settings. The directionality of the trajectory inferred by PAGA Tree was confirmed to be consistent with the RNA velocity–derived flow.

Trajectory inference was performed using Palantir (v1.3.6) on the literature dataset previously processed. Diffusion map was computed using the top principal components, and MAGIC was applied for data imputation. A highly proliferative cell, identified by elevated expression of *Mki67* and *Top2a*, was selected as the start cell. Terminal cell states were inferred based on the diffusion embedding, and pseudotime was subsequently computed. Cells were assigned to fates based on terminal state probabilities, retaining those with >70% probability of belonging to a given fate for downstream DEG analysis. DEG between the two major fates were identified by comparing pseudotime-ordered cells using the Wilcoxon rank-sum test. Genes expressed in at least 10% of cells in one of the groups were retained, and significance thresholds were set at adjusted *p* < 0.05 and absolute log₂ fold change > 0.58. Fate-specific upregulated genes were subjected to pathway enrichment analysis using Enrichr against the WikiPathways 2019 Mouse database. Enriched terms were compared between fates to highlight divergent biological programs.

### GRN inference

Gene regulatory networks were inferred using the SCENIC pipeline, applied to ASC (wild-type) populations from non-injured (NI) and 14-hour post-injury (Inj_14h) conditions, on a high-performance computing cluster. SCENIC was run with default parameters, including the use of pySCENIC modules for co-expression network inference, motif enrichment, and regulon activity scoring. Downstream analysis was restricted to Limch1⁺ ASCs from non-injured (NI) and 14-hour post-injury (Inj_14h) conditions. Differential regulon specificity between conditions was quantified using the ΔRSS metric. Regulons with |ΔRSS| > 0.05 and ≤300 target genes were retained, and those upregulated in either NI or Inj_14h were visualized. Regulon activity scores were compared between conditions using the Wilcoxon rank-sum test. Selected regulons of interest were visualized using violin plots to illustrate shifts in activity across conditions.

### ASC and FAP classification

To identify activated stromal cells (ASCs) among fibro-adipogenic progenitors (FAPs) in injured muscle, we trained logistic regression classifiers on single-cell transcriptomic profiles from injured KikGR mice. As the ASC labeling relies on KikGR gene expression labeling, which doesn’t capture all migratory ASCs, the classification task was approached with caution due to label noise and substantial class imbalance (fewer ASC-labeled cells than FAPs).

Feature selection was first performed using L1-penalized logistic regression (Lasso), which retained a sparse set of informative genes. A grid search over hyperparameters (regularization strength, solver, iteration limit, class weighting) and undersampling strategies was then performed, using stratified fivefold cross-validation. To account for the rarity of ASCs and the imperfect labeling inherent to the KikGR system, two classification objectives were evaluated:

- A balanced model, selected to maximize the minimum recall between ASC and FAP predictions, ensuring reliable detection of both populations.
- A high-sensitivity model, chosen to maximize ASC recall, with secondary sorting on FAP recall to avoid overfitting.

The classifier was trained on the Lasso-selected gene set and evaluated on a held-out test set. Post hoc threshold adjustment was applied to reduce ASC overcalling among FAPs by reverting low-confidence ASC predictions (probability < 0.7) back to FAP. Downstream analysis was done on the high-sensitivity model classification.

Model interpretability was further addressed using SHAP (SHapley Additive exPlanations). SHAP values were computed for each gene to assess their contribution to ASC classification. These values were compared with model coefficients to highlight key discriminants genes, revealing a small set of genes with high predictive and explanatory power.

The signature used on ASC injured conditions was based on top 10 predictive genes from the model coefficients for ASC (positive) and FAP (negative). Top 20% ASC with highest score were labelled as “High positive” cells for this signature.

## Data Availability

Processed data generated in this study will be deposited in a public repository upon acceptance of the article. The publicly available scRNA-seq datas re-analyzed in this study were obtained through the GEO database under accession codes: <u>GSE143437</u> and <u>GSE232106</u>. Data generated in this study is submitted to GEO under accession codes GSE307346 and GSE307346 and will be available upon publication of this article.

## Code Availability

Scripts used for the analyses presented in this study are available on GitHub (https://github.com/Nagooz).

Some illustrations were created with BioRender.com.

## Acknowledgements

We are grateful to the RESTORE CERT platforms, Mathieu Vignau for imaging analysis. Microscopy, RNA and library preparations were performed on equipment belonging to the RESTORE “Centre d’Expertise et de Ressources Technologiques“ UMR 1301-INSERM 5070-CNRS.

We acknowledge the animal facility US006/ CREFRE/INSERM/UPS.

We thank Emeline Sarot and Carine Valle from the sequencing platform, CRCT Technological cluster (INSERM-UMR1037) for support and assistance. We are grateful to Juan Pablo Cerapio Arroyo and Marion Perrier for assistance and advice (bioinformatics platform, CRCT Technological cluster (INSERM-UMR1037)).

This work was financially supported by INSERM, CNRS, Etablissement Français du Sang (EFS), ANR 20-CE19-0010 (CS), AFM-Téléthon PhD grant 23384 (MM) and Trampolin Grant 23263 (AG), ANR JCJC-21-CE14-0020 (AG), and through the grant EUR CARe N°ANR-18-EURE-0003 in the framework of the Programme des Investissements d’Avenir (ML).

## Supplementary data

**Supplementary Fig.1:**
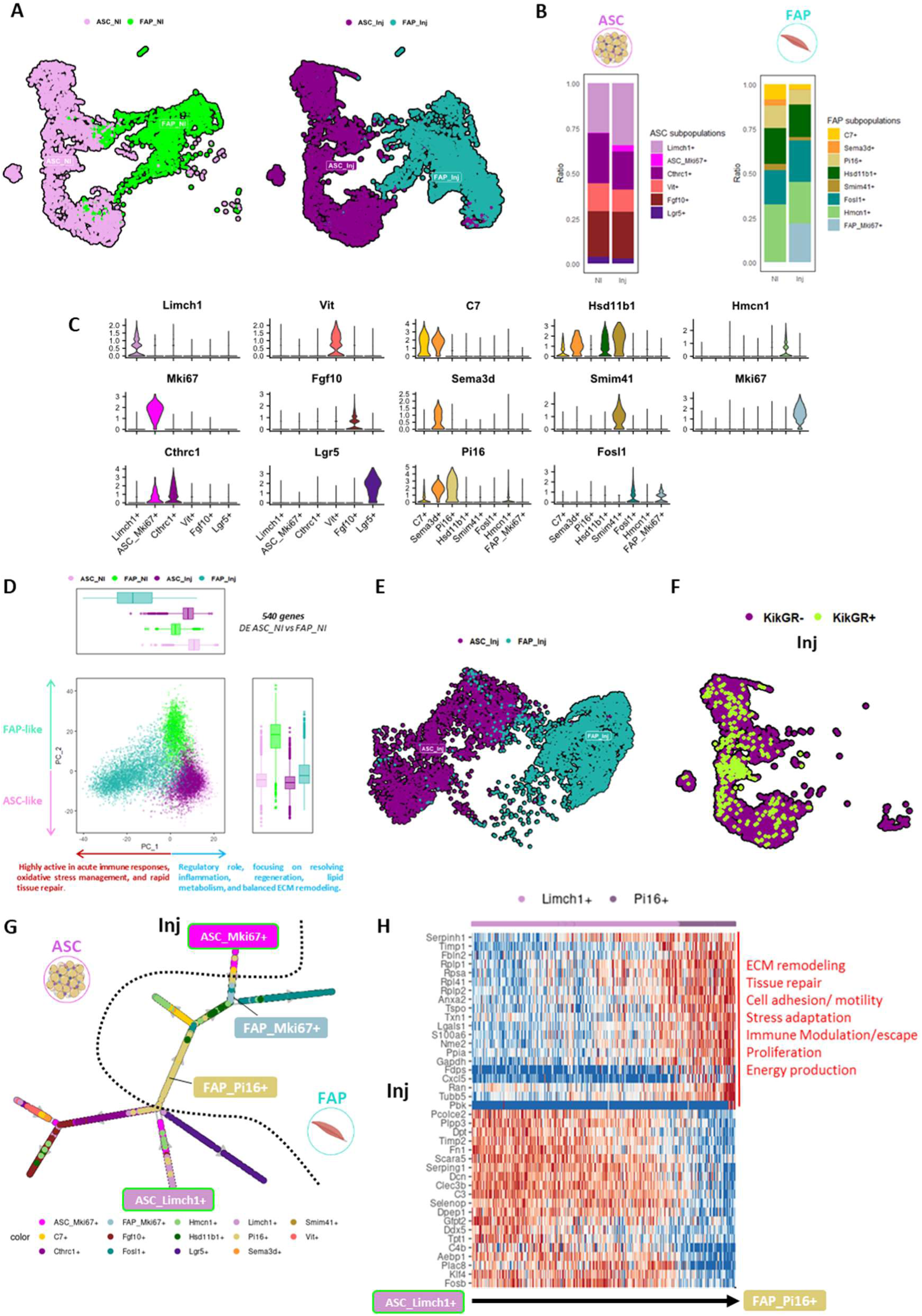
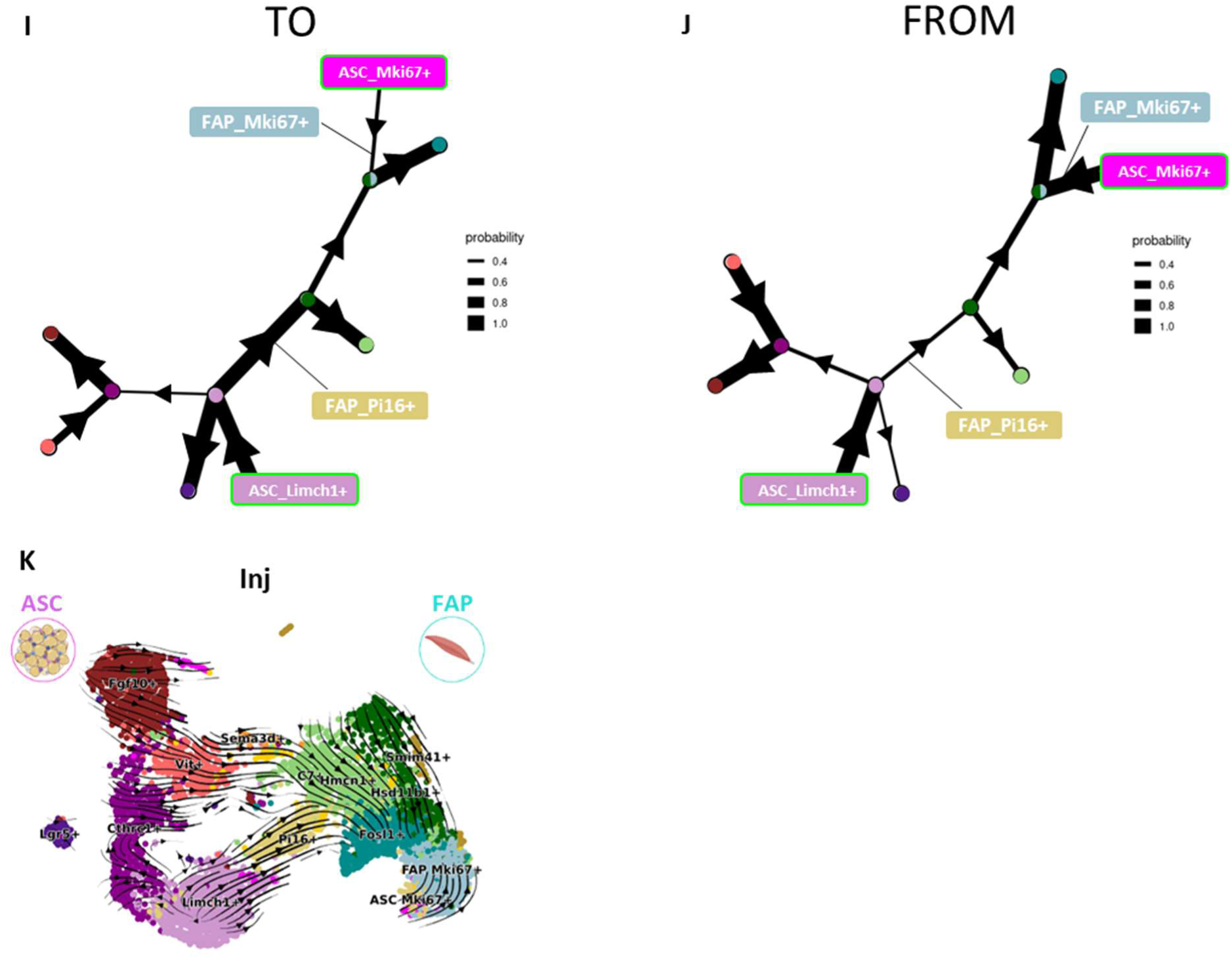
Progenitor and Proliferative ASC subpopulations from the scAT contribute to FAP heterogeneity in muscle regeneration. **A** UMAP embedding of ASCs and FAPs scRNA-seq data split by non-injured (NI) and Injured (Inj) conditions, colored by cell type/condition combination. **B** Barplots of sub-populations composition of ASCs (left) and FAPs (right) across conditions. **C** Violin plots showing the expression levels of representative marker genes of each ASCs and FAPs sub-populations. **D** Principal component analysis (PCA) of ASCs and FAPs under NI and Inj conditions, projected onto the space defined by the 540 genes differentially expressed between nin-injured ASC and FAPs (ASC_NI and FAP_NI). Each point represents a cell, colored by cell type and condition. Boxplots on the axes show the distribution of each cell type/condition combination along the principal components. **E** UMAP projection of ASCs and FAPs in the injured condition (ASC_Inj and FAP_Inj) onto the space defined by the 540 genes differentially expressed between ASCs and FAPs in the non-injured condition (ASC_NI and FAP_NI). This visualization highlights the transcriptional relationship between ASC and FAP populations in the injured condition, within a gene space informed by their baseline differences. **F** UMAP projection of ASCs in the injured condition (ASC_Inj), with KikGR+ cells highlighted. **G** 2D plot of the trajectory inference using PAGA Tree of the ASCs and FAPs sub-populations in the injredcondition (Inj), cells are colored by clusters. Dotted line separate ASCs (left) from FAPs (right). **H** Heatmap of top 40 differentially expressed genes through the trajectory ASC_Limch1^+^ and FAP_Pi16^+^ during regeneration, to show their dynamic expression along pseudotime. Gene Ontology terms linked to the genes up-regulated through the trajectory are visible in red. **I-J** 2D plot of the trajectory inference using PAGA Tree of the ASCs and FAPs sub-populations in the injured condition (Inj), showing the probability for a node to give rise to (**I**) or come from (**I**) a certain node. Nodes are colored based on their main sub-population composition. **K** RNA velocity analysis of ASCs and FAPs under the injured condition (Inj), performed using UniTVelo. Arrows indicate the inferred direction and magnitude of transcriptional dynamics, reflecting the predicted future states of cells.

**Supplementary Fig.2:**
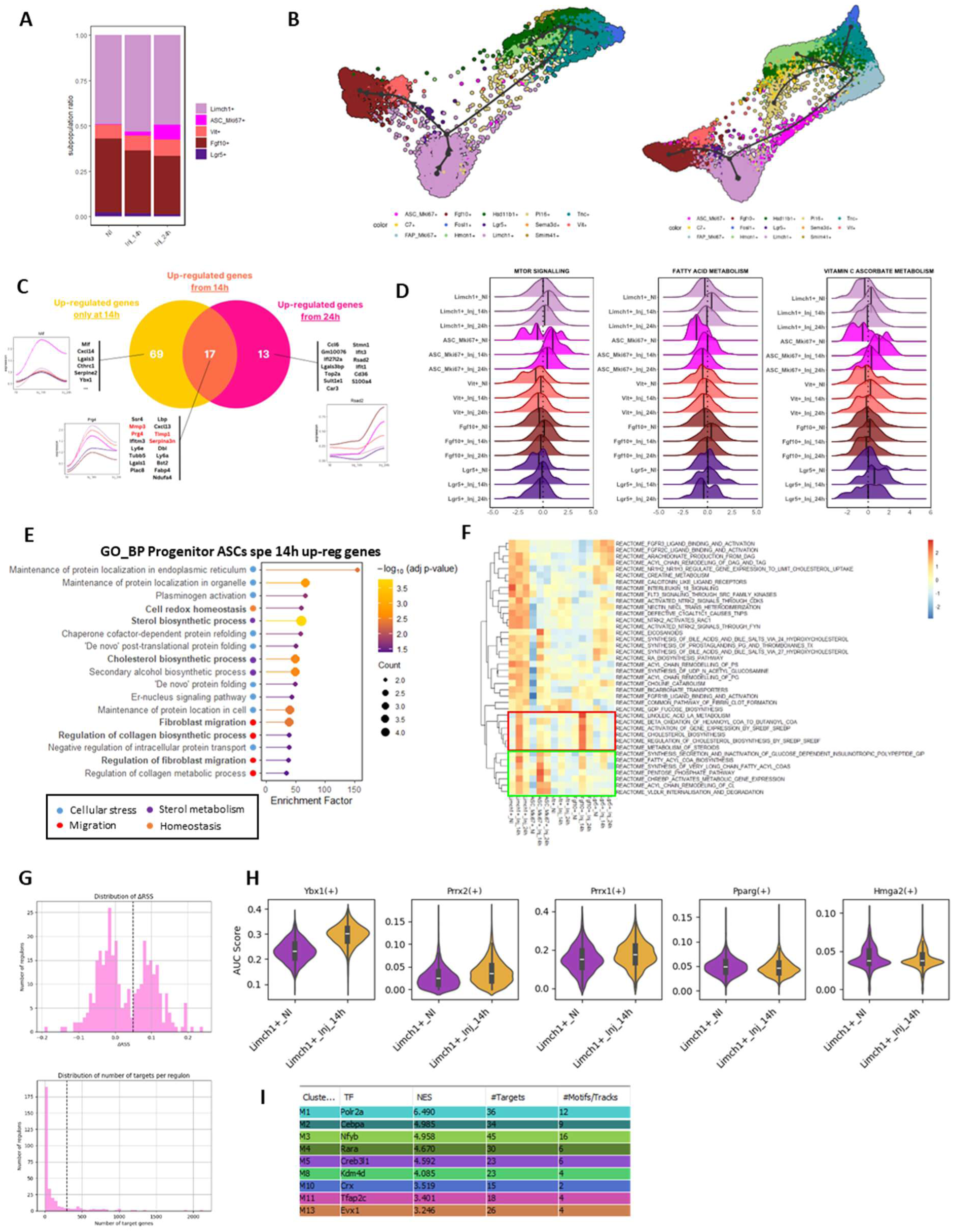
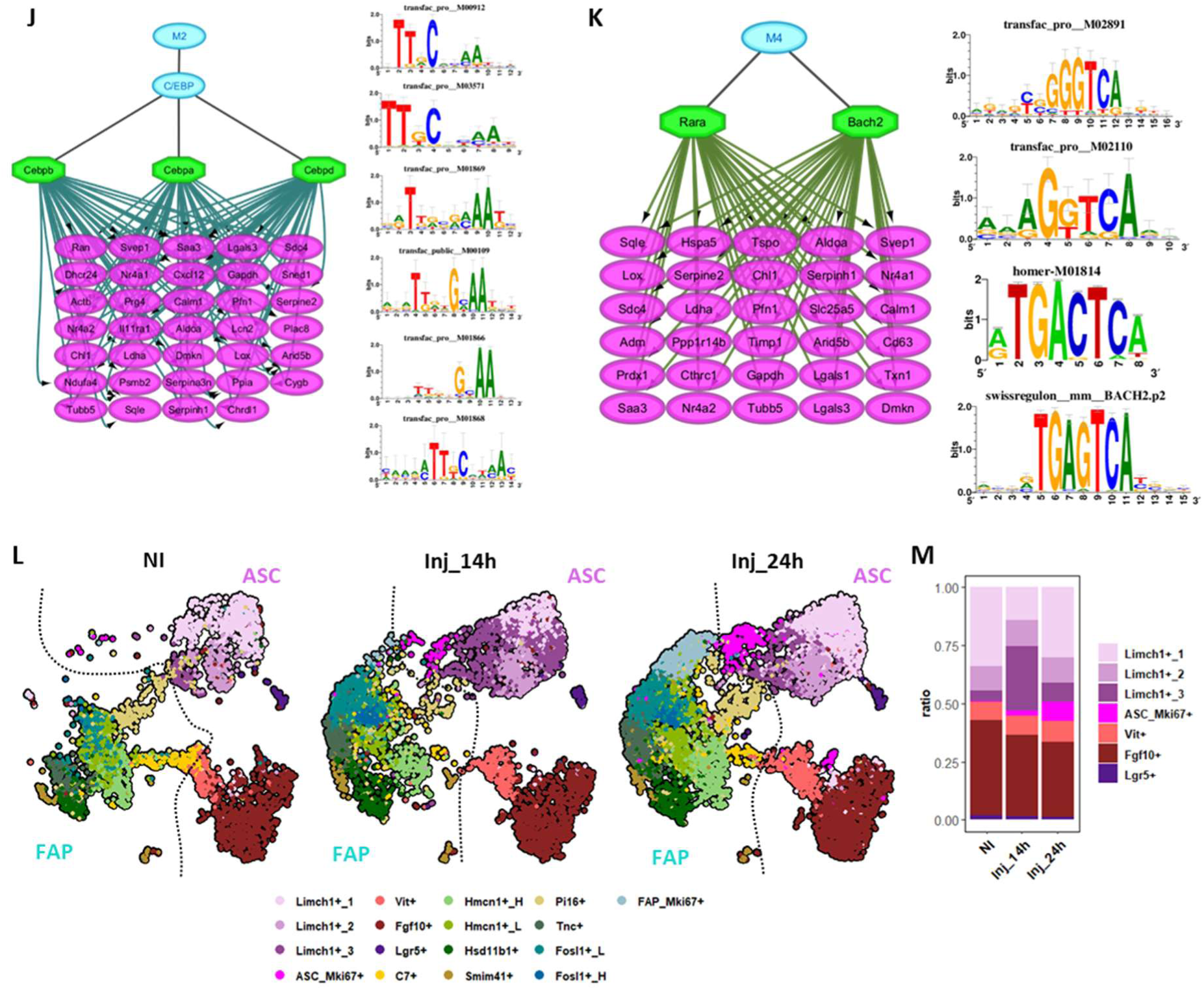
Early activation of migrating Progenitor ASCs with regenerative and adipogenic potential. **A** Barplots of sub-populations composition of ASCs across conditions. **B** Trajectory inference using PAGA Tree (Partition-based graph abstraction) of the ASC and FAP sub-populations at 24h post-injury (Inj_24h). **C** Venn diagram showing overlap of DEGs in ASCs in the injured condition at 14 and 24h (Inj_14h and Inj_24h) compared to the non-injured (NI) condition. Line plots illustrate representative gene expression kinetics for each category. **D** Ridge plots showing the distribution of selected Reactome pathway scores across ASC subpopulations and conditions. Selected pathways are upregulated in ASC Progenitors following injury. **E** Pathway enrichment analysis of specifically upregulated genes in ASC Progenitors at 14h post-injury. Gene Ontology (GO), KEGG, and Reactome databases were queried. Enriched terms related to cellular stress are marked with a blue dot, to migration to a red one, to sterol metabolism to a purple one and to homeostasis to an orange one. **F** Reactome enrichment analysis of upregulated genes in ASC sub-populations across conditions. Green box highlights pathways upregulated in ASC Progenitors at 14h post-injury (Inj_14h); red box indicates those also upregulated in ASC Preadipocytes at 14h post-injury (Inj_14h). Both sets of pathways are associated with sterol metabolism. **G** SCENIC regulon metrics. (Top) Distribution of regulons by ASC Progenitors ΔRSS (Inj_14h - NI) values. (Bottom) Distribution of regulons by number of target genes. **H** Violin plots showing AUC scores of selected regulons in ASC Progenitors under NI and Inj_14h conditions. **I** Transcription factors identified by iRegulon for genes upregulated in ASC Progenitors at 14h post-injury (Inj_14h) versus the non-injured (NI) condition. The table lists the normalized enrichment score (NES), number of predicted target genes, and number of associated motifs for each transcription factor. **J-K** Gene regulatory networks for transcription factors associated with specific modules. (J) C/EBP regulon linked to module M2. (K) Rara and Bach2 regulons linked to module M4. Right panels show the corresponding DNA-binding motifs for each transcription factor. **L** UMAP embedding of ASCs and FAPs split by non-injured (NI), 14h (Inj_14h) and 24h (Inj_24h) post-injury conditions, colored by clusters and subclusters for ASC Progenitors. Ten FAP sub-populations were identified using SNN (Shared Nearest Neighbor) clustering applied to each cell type. Dotted line separate ASCs (left) from FAPs (right). **M** Barplots of sub-populations composition of ASCs, with ASC Progenitors subclusters, across conditions.

**Supplementary Fig.3:**
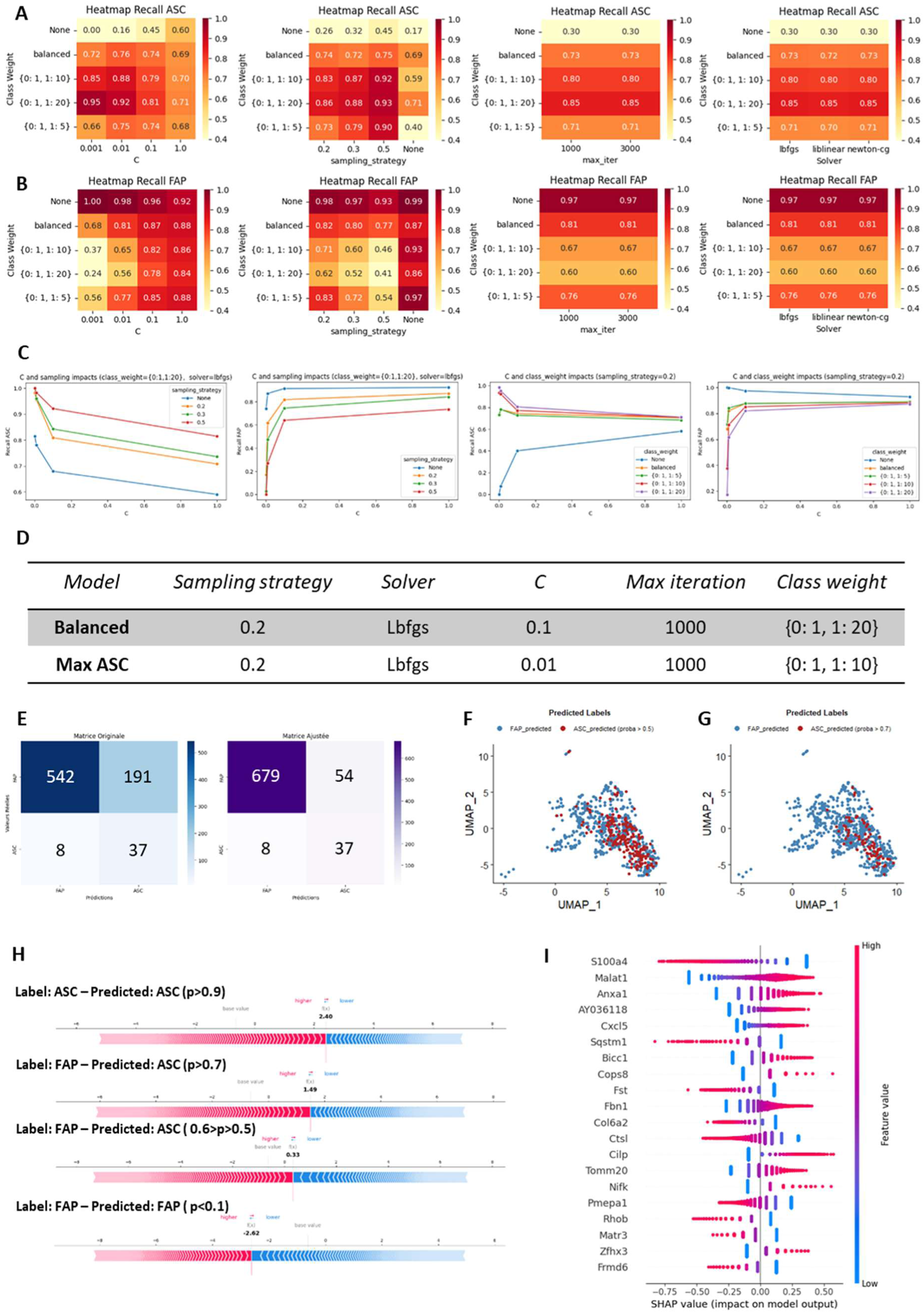
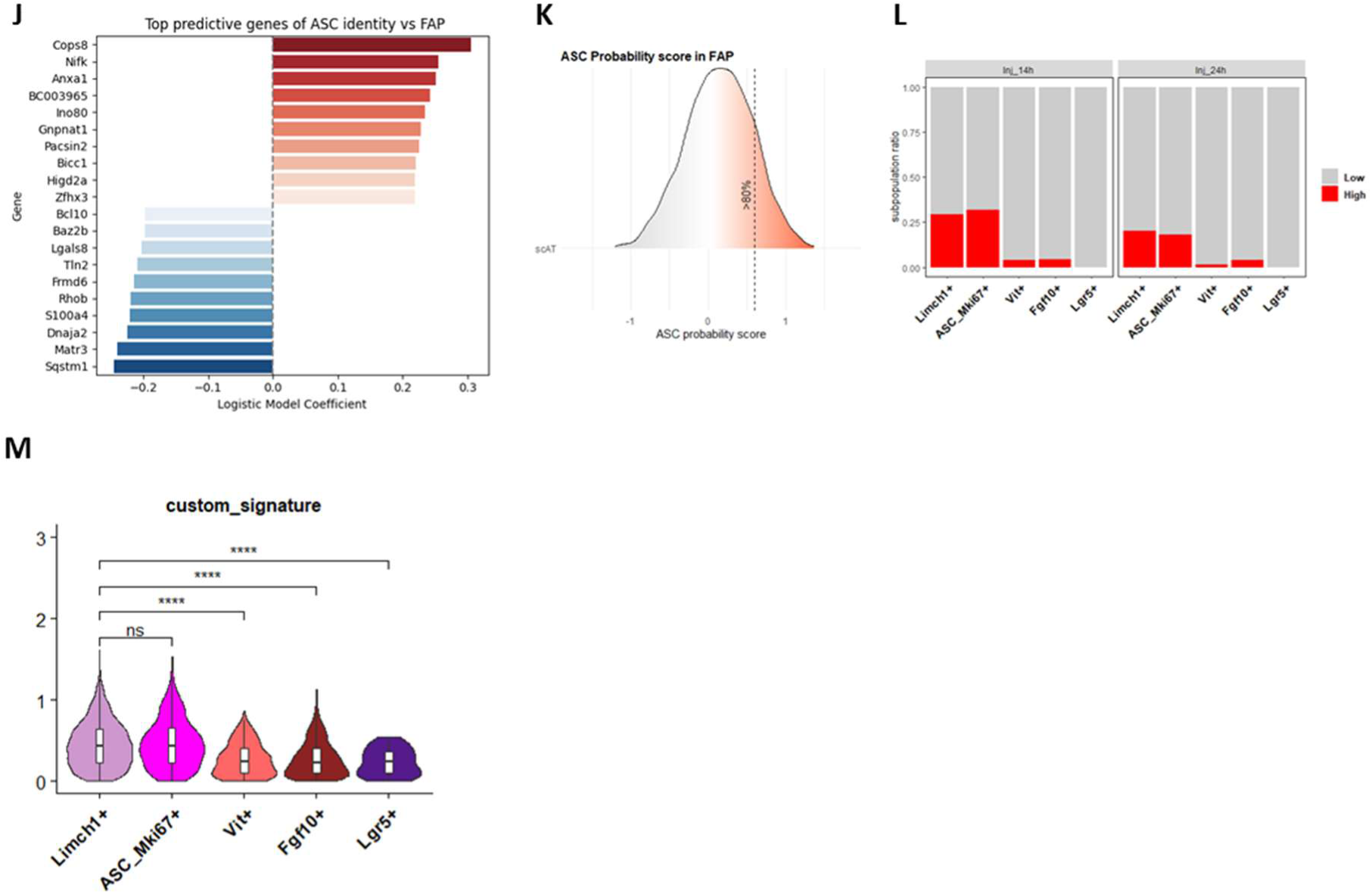
Logistic regression identifies a distinct Progenitor ASCs signature among FAP for tracing and functional prediction. **A-B** Heatmaps showing ASC (**A**) or FAP (**B**) recall across different logistic regression hyperparameter settings. Each panel varies one parameter, regularization strength (C), sampling strategy, number of iterations (max_iter), and solver, while displaying performance for multiple class weighting strategies. **C** Line plots showing the impact of regularization strength (C), sampling strategy, and class weight on classification performance. (Left two panels) ASC and FAP recall for different sampling strategies at fixed class weight and solver. (Right two panels) ASC and FAP recall for different class weight configurations at fixed sampling strategy. All models use logistic regression with solver = lbfgs. **D** Table summarizing the two logistic regression models selected, their hyperparameters and sampling strategy. **E** Confusion matrices showing model predictions for FAP and ASC classification. (Left) “Max ASC” model. (Right) “Max ASC” corrected model. **F-G** UMAP projection of FAP_Inj cells, with cells predicted as ASC in red with p>0.5 (**F**) or p>0.7 (**G**) and those predicted as FAP in blue by “Max ASC” model. No KikGR+ cells are present in this subset. **H** SHAP force plots showing feature contributions to ASC vs FAP classification in FAP cells for “Max ASC” model. Each panel represents a subgroup based on predicted ASC probability: high-confidence ASC-like (p > 0.7), intermediate (0.5 < p ≤ 0.6), and confident FAP (p < 0.1). Red features push the prediction toward ASC identity, while blue features push it toward FAP. The score reflects the model output for each cell, calculated as the sum of the base value and the SHAP contributions; higher scores indicate stronger ASC predictions. **I** Top features and their SHAP value for the “Max ASC” model. **J** Top features predictive of ASC identity vs FAP and their coefficient for “Max ASC” model. **K** Ridge plot showing the distribution of ASCs across ASC-derived FAP signature scores. The grey vertical line marks the threshold corresponding to the top 20% of cells with the highest scores. **L** Bar plots showing the proportion of cells with low (grey) or high (red) ASC-derived FAP signature scores across ASC subpopulations at 14 or 24h post-injury (Inj_14h and Inj_24h). **M** Violin plots showing ASC-derived FAP signature scores across ASC subpopulations at 14h post-injury (Inj_14h). Statistical comparisons were performed using two-sided Wilcoxon rank-sum tests. *p < 0.05, **p < 0.01, ***p < 0.001, ***p < 0.0001, ns: p>0.05.

**Supplementary Fig.4:**
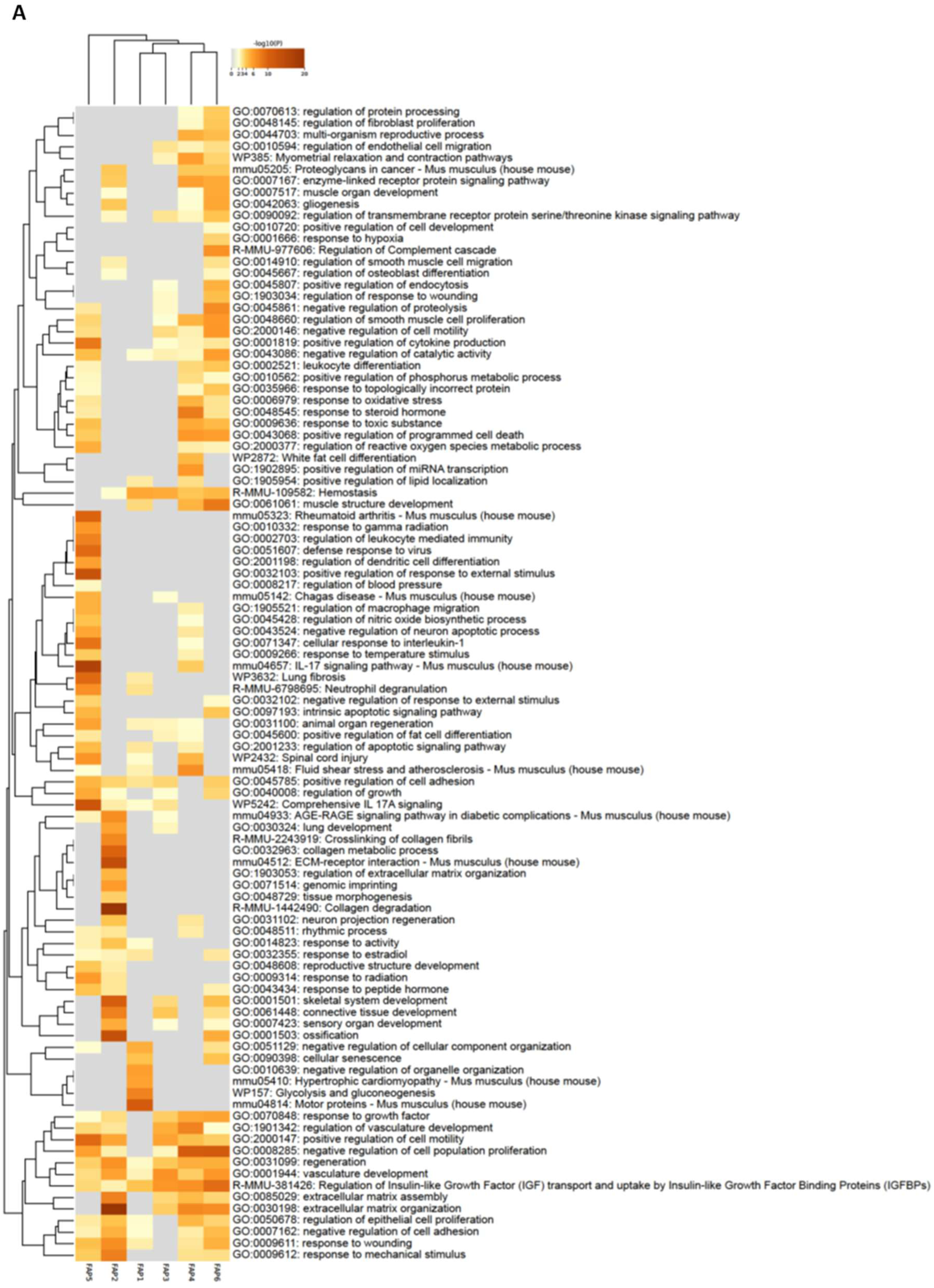
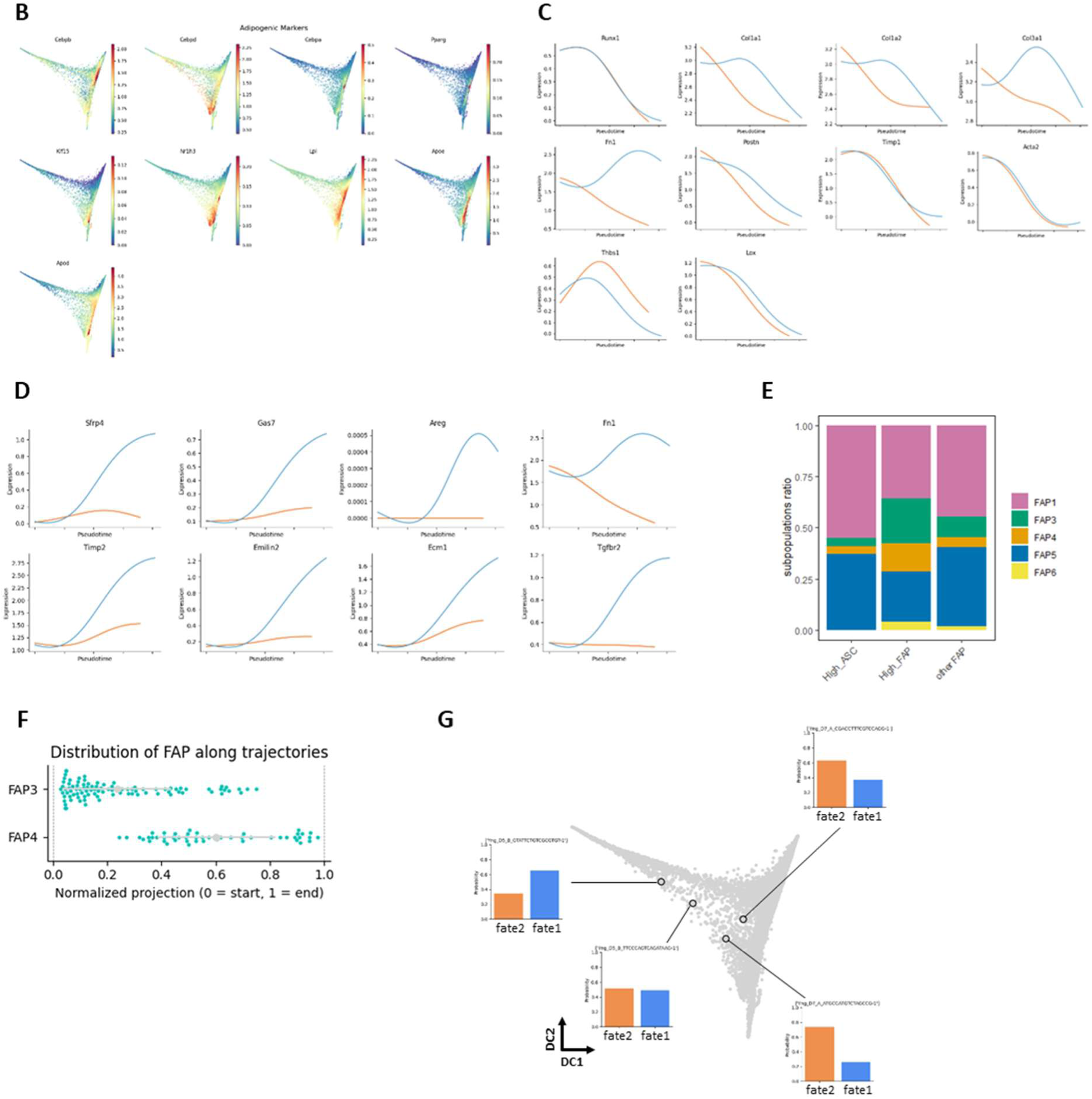
Progenitor ASCs among FAP exhibit an adipogenic bias along divergent regenerative trajectories. **A** Pathway enrichment analysis of Walter et al. FAP clusters. Gene Ontology (GO), KEGG, and Reactome databases were queried. **B** Feature plot of MAGIC treated key adipogenesis-related genes. **C** Expression dynamics of key fibrogenic-related genes along pseudotime for both fate1 (blue) and fate2 (orange) trajectories. **D** Expression dynamics of fate1 specific genes along pseudotime for both fate1 (blue) and fate2 (orange) trajectories. **E** Bar plots showing the composition in FAP clusters split by ASC-predicted FAPs and endogenous FAPs. **F** Scatter plot of the normalized distribution of predicted FAPs along the two differentiation trajectories represented by FAP3 and FAP4. **G** Diffusion map embedding of FAPs with bar plots showing the cell fate probabilities of some selected cells.

**Supplementary Fig.6:**
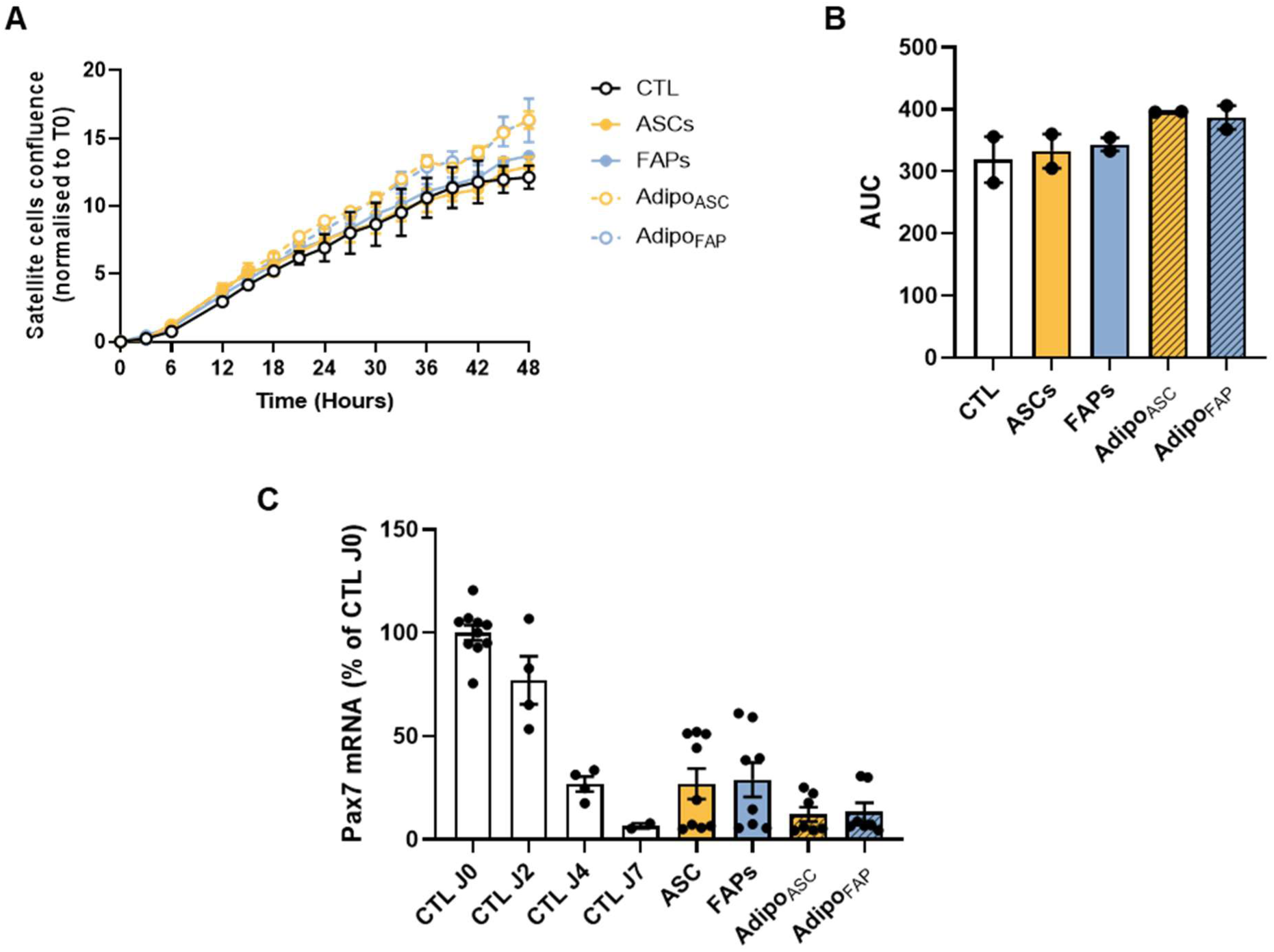
Adipocytes derived from ASCs are pro-myogenic. **A** Muscle stem cell (MuSC) proliferation assay performed with Incucyte. **B** Corresponding area under the curve (AUC). **C** Pax7 gene expression in differentiating MuSC cultivated with the several conditionned media for 4 days; control conditions without CM correspond to white bars. Results are expressed as mean ± SEM of 2 independant experiments; *p < 0.05.

**Supplementary Table S1:**
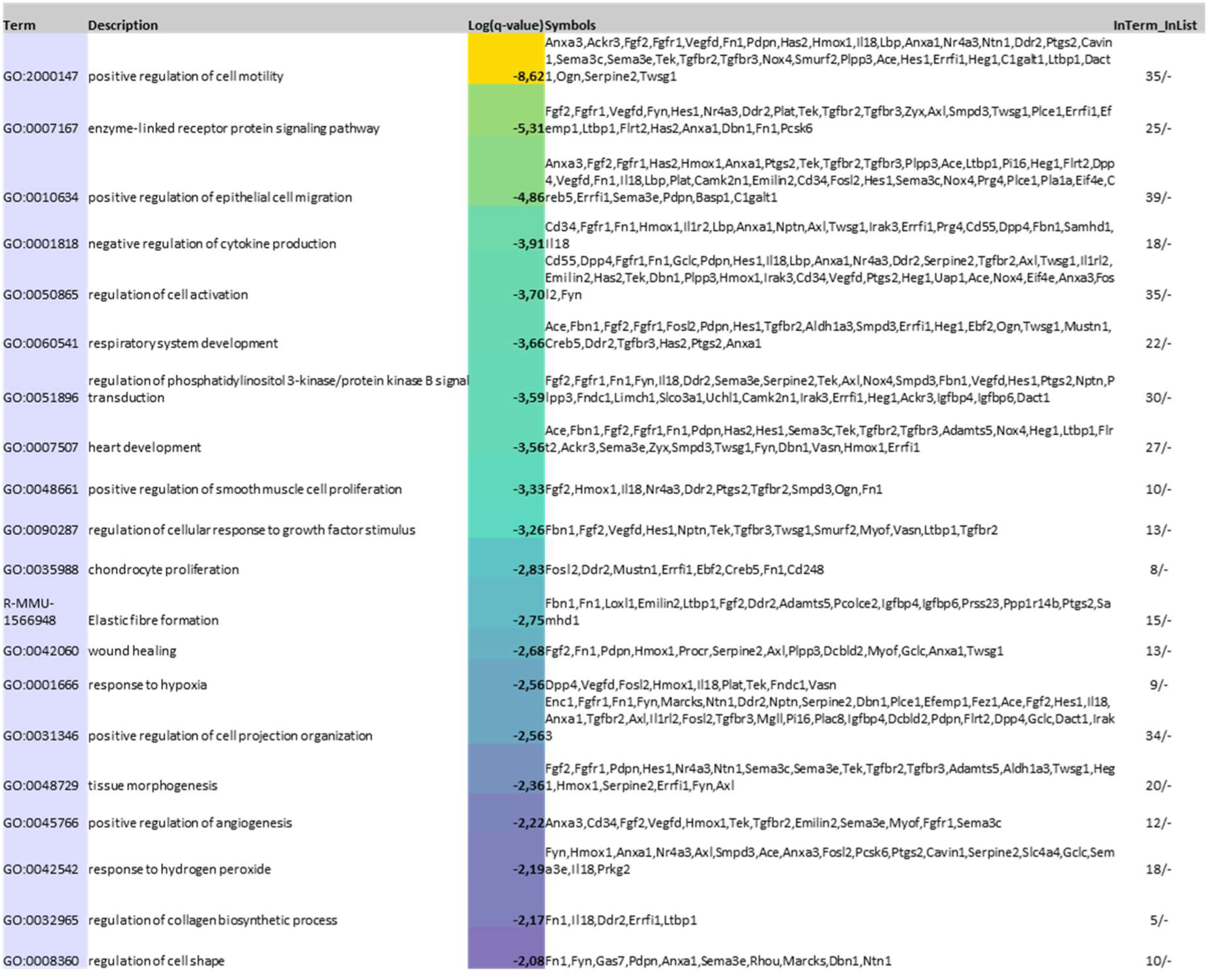
List of top 20 upregulated GO terms in Progenitor ASCs vs other ASCs in lnj.

**Supplementary Table S2.**
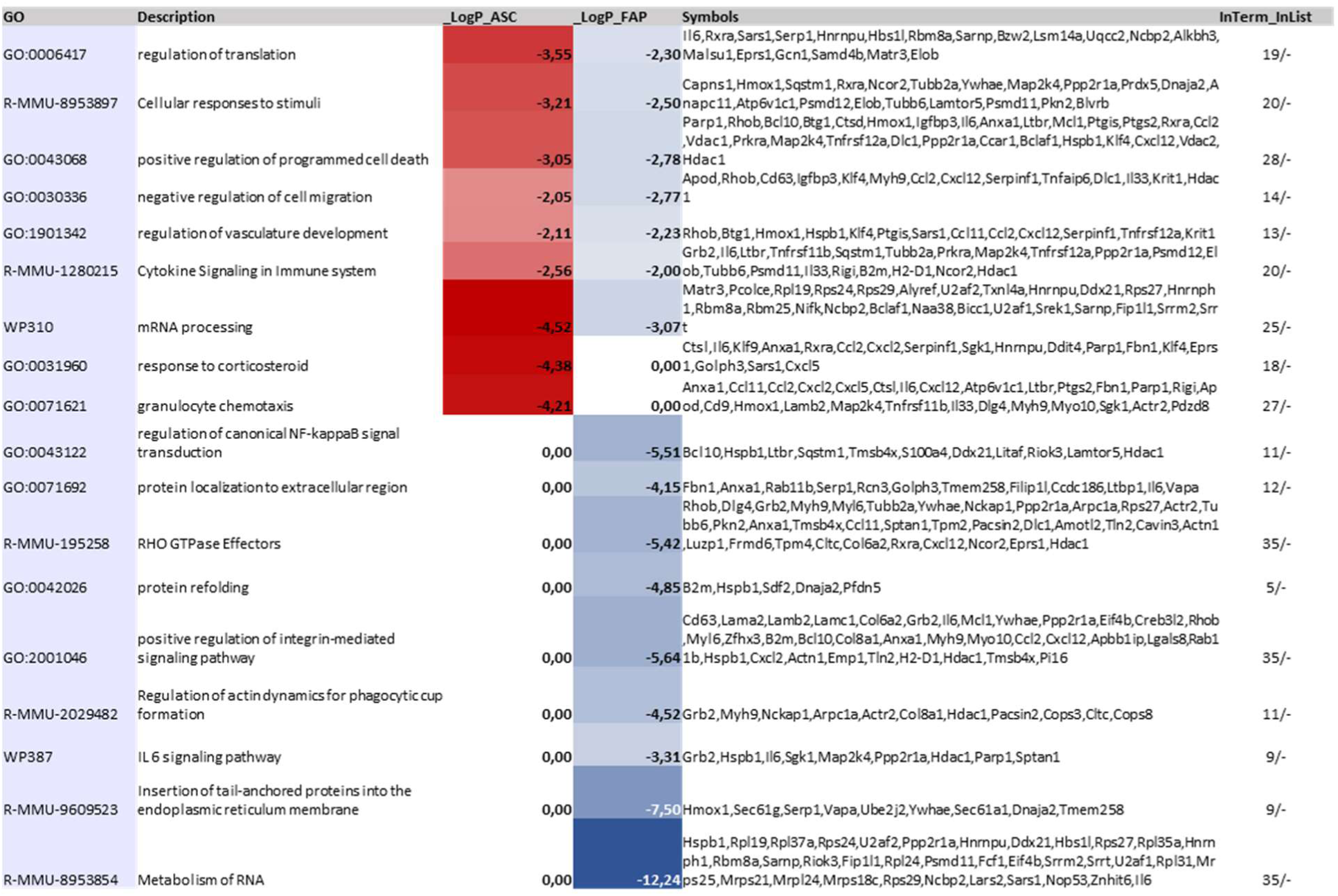
: **List of top upregulated GO terms in ASC- and FAP-associated gene signatures**

**Supplementary Table S3.**
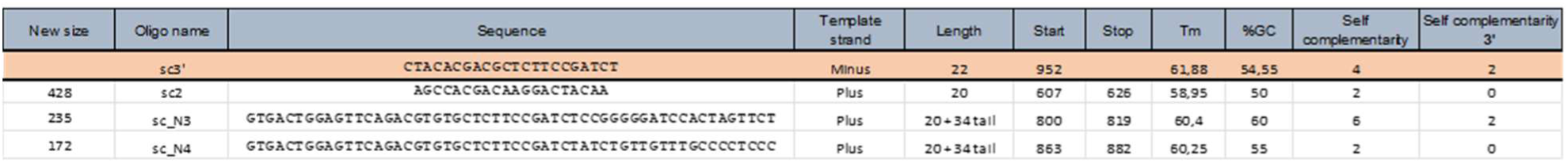
: **List of primers used for KikGR amplification by nested PCR**

## REFERENCES

1. Forcina, L., Cosentino, M. & Musarò, A. Mechanisms Regulating Muscle Regeneration: Insights into the Interrelated and Time-Dependent Phases of Tissue Healing. Cells 9, 1297 (2020).

2. Muire, P. J., Mangum, L. H. & Wenke, J. C. Time Course of Immune Response and Immunomodulation During Normal and Delayed Healing of Musculoskeletal Wounds. Front. Immunol. 11, 1056 (2020).

3. Joe, A. W. B. et al. Muscle injury activates resident fibro/adipogenic progenitors that facilitate myogenesis. Nat. Cell Biol. 12, 153–163 (2010).

4. Uezumi, A., Fukada, S., Yamamoto, N., Takeda, S. & Tsuchida, K. Mesenchymal progenitors distinct from satellite cells contribute to ectopic fat cell formation in skeletal muscle. Nat. Cell Biol. 12, 143–152 (2010).

5. Wosczyna, M. N., Biswas, A. A., Cogswell, C. A. & Goldhamer, D. J. Multipotent progenitors resident in the skeletal muscle interstitium exhibit robust BMP-dependent osteogenic activity and mediate heterotopic ossification. J. Bone Miner. Res. 27, 1004– 1017 (2012).

6. Theret, M., Rossi, F. M. V. & Contreras, O. Evolving Roles of Muscle-Resident Fibro-Adipogenic Progenitors in Health, Regeneration, Neuromuscular Disorders, and Aging. Front. Physiol. 12, 673404 (2021).

7. Oprescu, S. N., Yue, F., Qiu, J., Brito, L. F. & Kuang, S. Temporal Dynamics and Heterogeneity of Cell Populations during Skeletal Muscle Regeneration. iScience 23, 100993 (2020).

8. Rubenstein, A. B. et al. Single-cell transcriptional profiles in human skeletal muscle. Sci. Rep. 10, 229 (2020).

9. Malecova, B. et al. Dynamics of cellular states of fibro-adipogenic progenitors during myogenesis and muscular dystrophy. Nat. Commun. 9, 3670 (2018).

10. Fiore, D. et al. Pharmacological blockage of fibro/adipogenic progenitor expansion and suppression of regenerative fibrogenesis is associated with impaired skeletal muscle regeneration. Stem Cell Res. 17, 161–169 (2016).

11. Mathew, S. J. et al. Connective tissue fibroblasts and Tcf4 regulate myogenesis. Development 138, 371–384 (2011).

12. Uezumi, A. et al. Identification and characterization of PDGFRα+ mesenchymal progenitors in human skeletal muscle. Cell Death Dis. 5, e1186–e1186 (2014).

13. Murphy, M. M., Lawson, J. A., Mathew, S. J., Hutcheson, D. A. & Kardon, G. Satellite cells, connective tissue fibroblasts and their interactions are crucial for muscle regeneration. Development 138, 3625–3637 (2011).

14. Wosczyna, M. N. et al. Mesenchymal Stromal Cells Are Required for Regeneration and Homeostatic Maintenance of Skeletal Muscle. Cell Rep. 27, 2029–2035.e5 (2019).

15. Uezumi, A. et al. Mesenchymal *Bmp3b* expression maintains skeletal muscle integrity and decreases in age-related sarcopenia. J. Clin. Invest. 131, (2021).

16. Lemos, D. R. et al. Nilotinib reduces muscle fibrosis in chronic muscle injury by promoting TNF-mediated apoptosis of fibro/adipogenic progenitors. Nat. Med. 21, 786– 794 (2015).

17. Scott, R. W., Arostegui, M., Schweitzer, R., Rossi, F. M. V. & Underhill, T. M. Hic1 defines quiescent mesenchymal progenitor subpopulations with distinct functions and fates in skeletal muscle regeneration. Cell Stem Cell 25, 797–813.e9 (2019).

18. Contreras, O. et al. Cross-talk between TGF-β and PDGFRα signaling pathways regulates the fate of stromal fibro–adipogenic progenitors. J. Cell Sci. 132, jcs232157 (2019).

19. Dulauroy, S., Di Carlo, S. E., Langa, F., Eberl, G. & Peduto, L. Lineage tracing and genetic ablation of ADAM12(+) perivascular cells identify a major source of profibrotic cells during acute tissue injury. Nat. Med. 18, 1262–1270 (2012).

20. Sastourné-Arrey, Q. et al. Adipose tissue is a source of regenerative cells that augment the repair of skeletal muscle after injury. Nat. Commun. 14, 80 (2023).

21. Gil-Ortega, M. et al. Native adipose stromal cells egress from adipose tissue in vivo: evidence during lymph node activation. Stem Cells Dayt. Ohio 31, 1309–1320 (2013).

22. Gil-Ortega, M., Fernández-Alfonso, M. S., Somoza, B., Casteilla, L. & Sengenès, C. Ex vivo microperfusion system of the adipose organ: a new approach to studying the mobilization of adipose cell populations. Int. J. Obes. 38, 1255–1262 (2014).

23. Girousse, A. et al. The Release of Adipose Stromal Cells from Subcutaneous Adipose Tissue Regulates Ectopic Intramuscular Adipocyte Deposition. Cell Rep. 27, 323–333.e5 (2019).

24. Merrick, D. et al. Identification of a mesenchymal progenitor cell hierarchy in adipose tissue. Science 364, eaav2501 (2019).

25. Emont, M. P. et al. A single-cell atlas of human and mouse white adipose tissue. Nature 603, 926–933 (2022).

26. Schwalie, P. C. et al. A stromal cell population that inhibits adipogenesis in mammalian fat depots. Nature 559, 103–108 (2018).

27. Das, N., Schmidt, T. A., Krawetz, R. J. & Dufour, A. Proteoglycan 4: From Mere Lubricant to Regulator of Tissue Homeostasis and Inflammation. BioEssays 41, 1800166 (2019).

28. Ninkovic, N. et al. Proteoglycan 4 (PRG4) treatment improves skin wound healing in a porcine model. FASEB J. 38, e23547 (2024).

29. Krawetz, R. J. et al. Proteoglycan 4 (PRG4) treatment enhances wound closure and tissue regeneration. NPJ Regen. Med. 7, 32 (2022).

30. Dong, Y., Silva, K. A. S., Dong, Y. & Zhang, L. Glucocorticoids increase adipocytes in muscle by affecting IL-4 regulated FAP activity. FASEB J. 28, 4123–4132 (2014).

31. Ayala-Sumuano, J.-T. et al. Glucocorticoid Paradoxically Recruits Adipose Progenitors and Impairs Lipid Homeostasis and Glucose Transport in Mature Adipocytes. Sci. Rep. 3, 2573 (2013).

32. Cerquone Perpetuini, A., et al. Janus effect of glucocorticoids on differentiation of muscle fibro/adipogenic progenitors. Sci. Rep. 10, 5363 (2020).

33. Walter, L. D. et al. Transcriptomic analysis of skeletal muscle regeneration across mouse lifespan identifies altered stem cell states. *Nat*. Aging 4, 1862–1881 (2024).

34. Mahdy, M. A. A. Glycerol-induced injury as a new model of muscle regeneration. Cell Tissue Res. 374, 233–241 (2018).

35. Arsic, N. et al. Vascular endothelial growth factor stimulates skeletal muscle regeneration in Vivo. Mol. Ther. 10, 844–854 (2004).

36. Mahdy, M. A. A., Lei, H. Y., Wakamatsu, J.-I., Hosaka, Y. Z. & Nishimura, T. Comparative study of muscle regeneration following cardiotoxin and glycerol injury. Ann. Anat. - Anat. Anz. 202, 18–27 (2015).

37. Pisani, D. F. et al. Isolation of a Highly Myogenic CD34-Negative Subset of Human Skeletal Muscle Cells Free of Adipogenic Potential. Stem Cells 28, 753–764 (2010).

38. Pisani, D. F., Bottema, C. D. K., Butori, C., Dani, C. & Dechesne, C. A. Mouse model of skeletal muscle adiposity: A glycerol treatment approach. Biochem. Biophys. Res. Commun. 396, 767–773 (2010).

39. Lukjanenko, L., Brachat, S., Pierrel, E., Lach-Trifilieff, E. & Feige, J. N. Genomic Profiling Reveals That Transient Adipogenic Activation Is a Hallmark of Mouse Models of Skeletal Muscle Regeneration. PLOS ONE 8, e71084 (2013).

40. Dammone, G. et al. PPARγ Controls Ectopic Adipogenesis and Cross-Talks with Myogenesis During Skeletal Muscle Regeneration. Int. J. Mol. Sci. 19, 2044 (2018).

41. Wagatsuma, A. Adipogenic potential can be activated during muscle regeneration. Mol. Cell. Biochem. 304, 25–33 (2007).

42. Kotsaris, G. et al. Odd skipped-related 1 controls the pro-regenerative response of fibro-adipogenic progenitors. Npj Regen. Med. 8, 19 (2023).

43. Arrighi, N. et al. Characterization of adipocytes derived from fibro/adipogenic progenitors resident in human skeletal muscle. Cell Death Dis. 6, e1733–e1733 (2015).

44. Norris, A. M. et al. Intramuscular adipose tissue restricts functional muscle recovery. Cell Rep. 44, 116021 (2025).

45. Flores-Opazo, M. et al. Fibro-adipogenic progenitors in physiological adipogenesis and intermuscular adipose tissue remodeling. Mol. Aspects Med. 97, 101277 (2024).

46. Norris, A. M. et al. Studying intramuscular fat deposition and muscle regeneration: insights from a comparative analysis of mouse strains, injury models, and sex differences. Skelet. Muscle 14, 12 (2024).

47. Zubiría, M. G. et al. Dexamethasone primes adipocyte precursor cells for differentiation by enhancing adipogenic competency. Life Sci. 261, 118363 (2020).

48. Mankodi, A. et al. Quantifying disease activity in fatty-infiltrated skeletal muscle by IDEAL-CPMG in Duchenne muscular dystrophy. Neuromuscul. Disord. NMD 26, 650– 658 (2016).

49. Hausman, G. J., Basu, U., Du, M., Fernyhough-Culver, M. & Dodson, M. V. Intermuscular and intramuscular adipose tissues: Bad vs. good adipose tissues. Adipocyte 3, 242–255 (2014).

50. Biltz, N. K. et al. Infiltration of intramuscular adipose tissue impairs skeletal muscle contraction. J. Physiol. 598, 2669–2683 (2020).

51. Liu, W., Liu, Y., Lai, X. & Kuang, S. Intramuscular adipose is derived from a non-Pax3 lineage and required for efficient regeneration of skeletal muscles. Dev. Biol. 361, 27–38 (2012).

52. Collins, K. H. et al. Leptin mediates the regulation of muscle mass and strength by adipose tissue. J. Physiol. 600, 3795–3817 (2022).

53. Lee, C. et al. Rotator Cuff Fibro-Adipogenic Progenitors Demonstrate Highest Concentration, Proliferative Capacity, and Adipogenic Potential Across Muscle Groups. J. Orthop. Res. 38, 1113–1121 (2020).

54. Duerre, D. J. & Galmozzi, A. Deconstructing Adipose Tissue Heterogeneity One Cell at a Time. Front. Endocrinol. 13, (2022).

55. Sárvári, A. K. et al. Plasticity of Epididymal Adipose Tissue in Response to Diet-Induced Obesity at Single-Nucleus Resolution. Cell Metab. 33, 437–453.e5 (2021).

56. Bäckdahl, J. et al. Spatial mapping reveals human adipocyte subpopulations with distinct sensitivities to insulin. Cell Metab. 33, 1869–1882.e6 (2021).

57. Massier, L. et al. An integrated single cell and spatial transcriptomic map of human white adipose tissue. Nat. Commun. 14, 1438 (2023).

58. Arrighi, N. et al. Characterization of adipocytes derived from fibro/adipogenic progenitors resident in human skeletal muscle. Cell Death Dis. 6, e1733–e1733 (2015).

59. Santoro, A., McGraw, T. E. & Kahn, B. B. Insulin action in adipocytes, adipose remodeling, and systemic effects. Cell Metab. 33, 748–757 (2021).

60. Martínez-Sánchez, N. There and Back Again: Leptin Actions in White Adipose Tissue. Int. J. Mol. Sci. 21, 6039 (2020).

61. Saelens, W., Cannoodt, R., Todorov, H. & Saeys, Y. A comparison of single-cell trajectory inference methods. Nat. Biotechnol. 37, 547–554 (2019).

